# Rational Design of Immunogenic Nanoparticles as a Platform for Enhanced Ovarian Cancer Immunotherapy in Mice

**DOI:** 10.1101/2025.06.05.657862

**Authors:** Lien Tang, Ben Marwedel, Caleb Dang, Marian Olewine, Melanie Jun, Paulina Naydenkov, Lorél Y. Medina, Veronica Gayoso, Ngoc Doan, Shamus L. O’Leary, Carmine Schiavone, Joseph Cave, Tamara Howard, John D. Watt, Prashant Dogra, Rita E. Serda, Achraf Noureddine

**Author notes:** equal contributions (Lien Tang, Ben Marwedel). equal contributions (Caleb Dang, Marian Olewine).

## Abstract

Ovarian cancer immunotherapy remains a challenge based on the “cold” tumor microenvironment. Herein we present a rational design to create immunogenic nanoparticles as a multi-agent platform that promotes immune response in a mouse model of ovarian cancer. The hybrid lipid-silica nanosystem is capable of co-loading four types of cargo molecules including a model antigen, nucleic acid-based adjuvant Cytosine-p-linked to Guanine (CpG, TLR3/9 agonist), lipid-based adjuvant (MPLA, TLR4 agonist) integrated into the lipid coat, and optionally a small molecule drug, such as the chemotherapeutic agent oxaliplatin, a well-established treatment for ovarian cancer. The optimization of the nanoplatform in terms of lipid composition, functionalized silica dendritic core formation, and final charge, as well as their compatibility with the complex loading profile highlights an opportunity for enhanced survival of mice with advanced ovarian cancer compared to monotherapy. Furthermore, intraperitoneal administration led to preferential accumulation within tumor-burdened tissues with selective accumulation in myeloid cells. High myeloid cell cytotoxicity negated the benefits of oxaliplatin. The inclusion of CpG in the nanoparticle formulation enhanced the survival of mice with ovarian cancer. To interpret these outcomes and guide future design, we also developed a mathematical model of nanoparticle-driven immune activation, which quantified treatment efficacy and identified key parameters governing tumor response. The presented hybrid nanoparticle is tunable, enabling delivery of alternative molecules therefore, thereby highlighting a promising platform for the treatment of peritoneal cancers.

**Graphical TOC:** 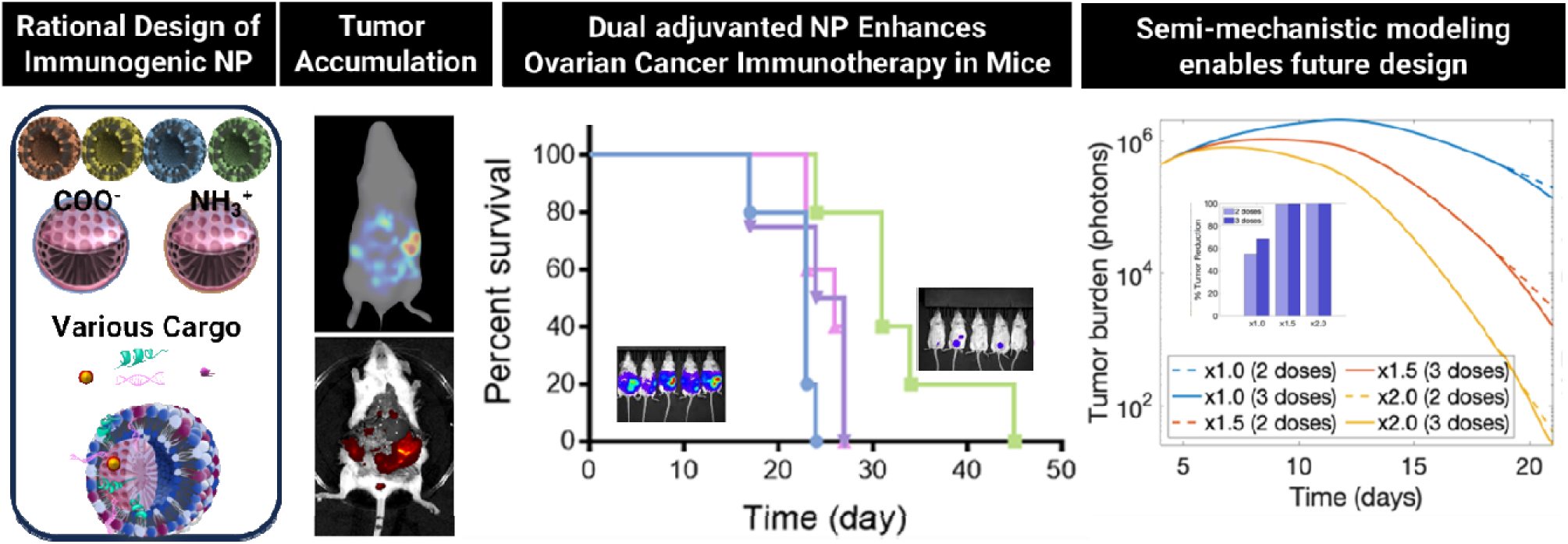

## 1. Introduction

The field of immunotherapy as an option for cancer treatment has grown immensely over the last several years. By stimulating the immune system to specifically target cancer cells, immunotherapy has shown to be an effective therapy to treat malignancy and prevent tumor recurrence while reducing systemic toxicity compared to conventional cancer treatment options^1–4^. One of the major limitations to the effectiveness of cancer immunotherapy is the influence of the tumor microenvironment, which is predominately immunosuppressive and hinders the ability of immune cells to function effectively^5, 6^. Nanomedicine and the ability of nanoparticles to act as delivery systems for adjuvants, cancer antigens, and other biomolecules, has led to the development of targeted therapies with enhanced efficacy and fewer side effects^7–9^. Facilitating the targeted delivery of cancer antigens or other desired cargo molecules to stimulate the immune system, nanoparticle-based immunotherapies represent a promising strategy to modulate immune responses and overcome the suppressive effects of the tumor microenvironment.

Combination therapy involving the co-delivery of multiple molecules of interest has gained increasing attention, as it promises to harness the advantages of several compounds spatially colocalized within a mother carrier. This approach allows for enhanced therapeutic outcomes and targeting through the simultaneous and eventually coordinated activity of molecules such as cancer antigens and immunomodulators in a single payload, not to mention the possibility of incorporating chemotherapeutics as well to induce immunogenic cell death (ICD)^10–12^. However, while multiple-compound loading offers unique opportunities, like the elegant multilayered polycaprolactone/polyethylene glycol/DOTAP polymer-lipid system^13^, a straightforward system optimized for a wide range of molecules remains a technically challenging task as it requires more complex features to accommodate the diversity in cargo types.

Here we present a strategy based on the lipid-silica hybrid nanosystem,^14–18^ to create an immunogenic lipid-MSN (ILM). The platform includes rationally designing inner and surface properties to maximize cargo presentation to sequentially stimulate anti-cancer immunity. The high loading capacity is driven by a high surface area, protection and diversity of cargo within, h the sponge-like porous core^19–21^ and a lipid shell that enables spatial co-localization of complementary molecules of different natures and sizes such as tumor antigens, oligonucleotide adjuvant, lipid adjuvant, and a small molecule drug.

In efforts to leverage nanoparticle immunotherapy against peritoneal cancers, we have recently shown that intraperitoneal administration of nanoparticles in a disseminated ovarian cancer model results in nanoparticle accumulation in tumor-burdened omentum, mesentery and fat pads. Localization within tumor-burdened tissues is agnostic to factors including presence of surface Pathogen Associated Molecular Pattern (PAMPs) or nanoparticle size (100 vs 500 nm)^22, 23^. This feature balances the limitations of intravenous injections in which nanoparticles accumulate predominately in liver and spleen. Localization in immune hotspots that lie within tumor-burdened tissues activates T cells in proximity to malignant cells.

Herein we present an engineered generic platform that considers three factors that bridge the remaining gaps in the particle design namely i) the lipid composition and its impact on the particles stability and homogeneity, ii) optimize the capability of multiple biomolecules loading while controlling the final charge of the nanoparticle, iii) yield mono vs dual adjuvanted nanoparticles with opposite charges each. Therefore, we have first carefully designed a library of eighteen lipid compositions to select optimal formulations that yield homogenous and stable nanoparticles, which next were leveraged to design a nanoparticles capable of co-loading complementary molecules, namely ovalbumin as a model antigen, nucleic acid- and lipid-based based adjuvants, and a chemotherapeutic, which shows macrophage activation, successful antigen processing and presentation, and enhanced survival of mice with ovarian cancer. In addition to the comprehensive investigation of these parameters, and in line with the recent FDA recommendations about promoting Novel Approaches Methodologies NAMs^24,25^, we have incorporated a semi-mechanistic modeling that couples tumor growth dynamics with innate immune activation to validate in vivo efficacy experiments and guide future design of nanoparticle vaccine.

## 2. Results and Discussion

### 2.1. Design of functional silica nanodendrites

Our group has refined the elegant synthesis of dendritic nanoparticles by Zhao^26–28^, to create straightforward and well established strategies to create silica nanodendrites using the biphasic stratification process^15, 29^ that allows for the fine tuning of their structure with various pores and morphologies where the reaction is initiated at the interface between the silica source containing organic phase and the surfactant and catalyst-rich aqueous phase (**Fig. 2A**). A current challenge is the anchoring of functional groups while preserving the defined structures and porosity. Traditionally, the *in situ* addition of the silica source along with the organosilane shows an acceptable degree of functionalization as both molecules will hydrolyze and condense at comparable rates. In this instance, the *in situ* addition helps creating more nucleating sites resulting in interconnected seeded particles with a large hydrodynamic size (**Fig. 2C,D**). Considering the slow activation and polymerization process in this reaction, we have adapted the delayed co-condensation method, initially introduced by Bein^30^ for MCM-41 type MSN to this new type of dendritic MSN, that first result in premade nanoparticles (overnight) followed by reaction the organosilane for a few hours to attach to the MSN surface, hence maintaining the nanoparticle dendritic structure (**Fig. 2A**). To determine the charge properties required to make immunogenic nanoparticles, dendritic MSNs with a pores size ∼ 7 nm were made following our previous reports. The structure was selected to support the loading of model antigen ovalbumin (OVA).

**Figure 1.**
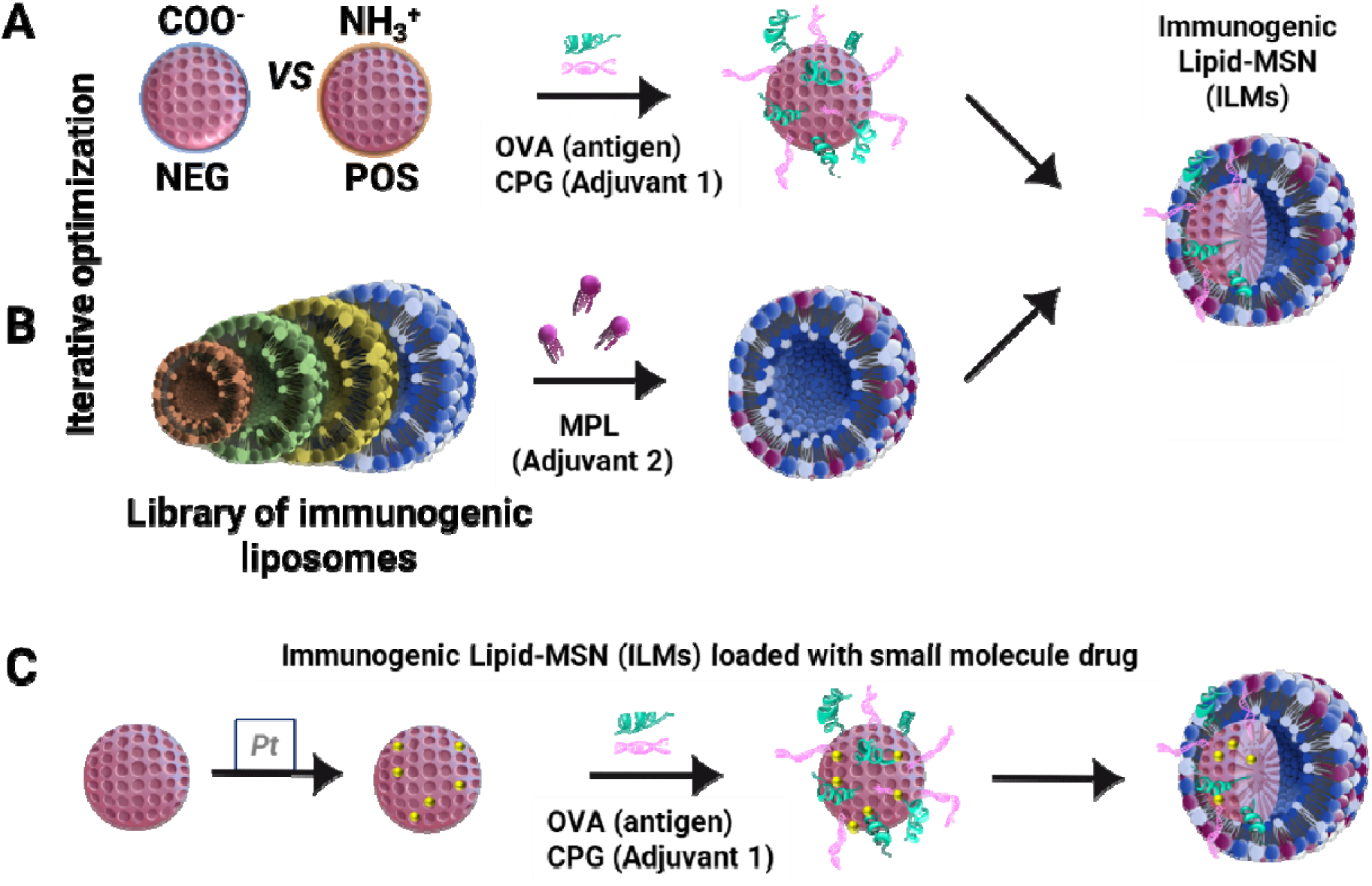
Schematic summarizing the design of the ILM platform. A) MSN with different surface chemistries resulting in different charges for optimal loading of OVA model antigen and CpG adjuvant, B) optimization of MPL-containing immunogenic lipid formulation

**Figure 2.**
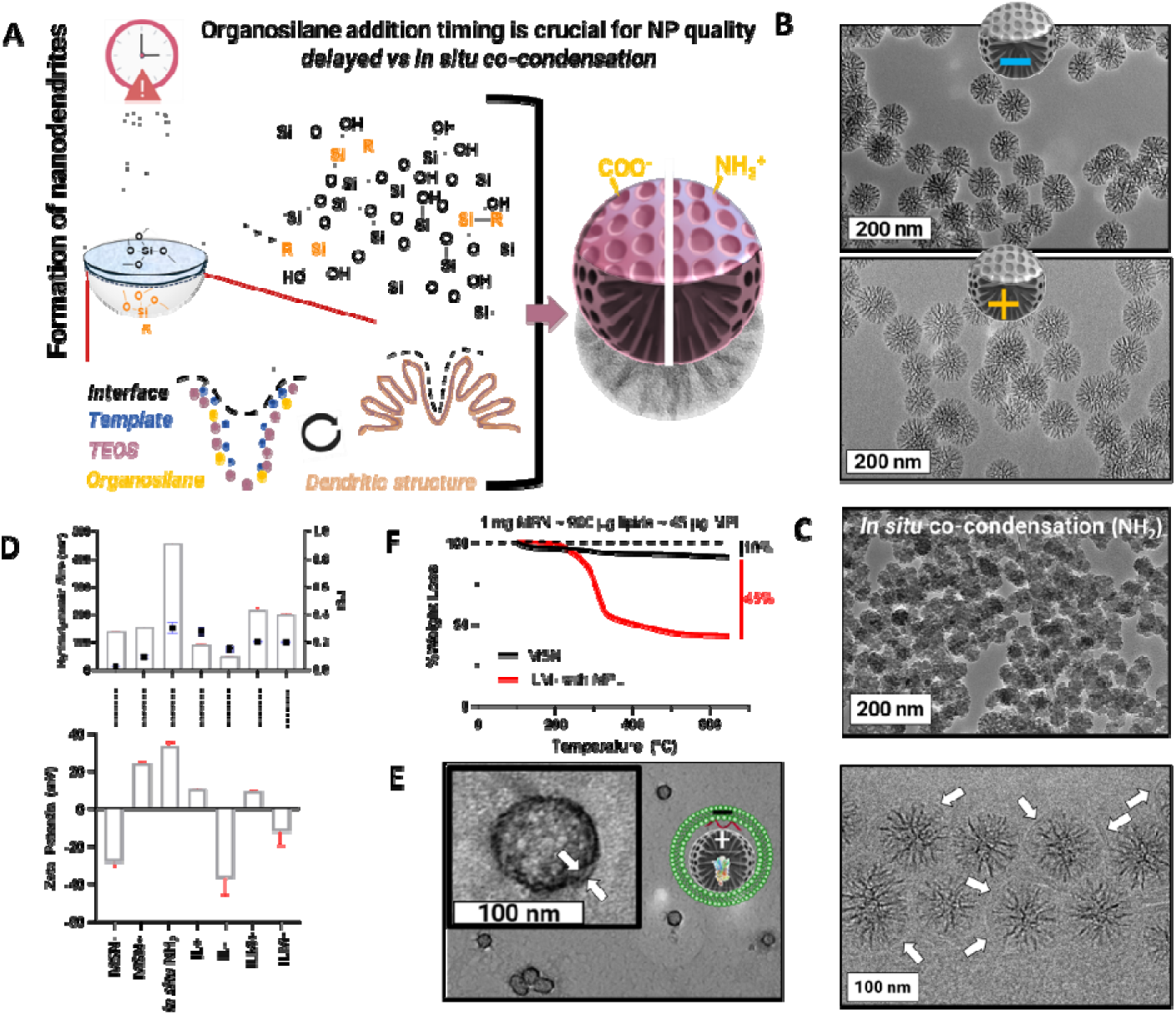
A) formation of the silica dendrite and crucial time of addition of organosilane B) TEM micrographs showing the dendritic structure of COOH- and NH_2_-functionalized MSN made via delayed co-condensation or C) in situ co-condensation, along with D) the respective size and zeta potential of the ILMs and their subcomponents, E) negatively stained TEM and CryoEM micrographs showing the bio-shell pointed out by white arrows F) TGA curves emphasizing the wt % of lipids and hence MPL in the nanoparticles.

TEM imaging showed monosized particles with a dendritic structure and ∼ 100 nm diameter (**Fig. 2B**) with a hydrodynamic size = 135 ± 2 nm and 150 ± 4 nm and zeta potential (ζ = -27 ± 4 and ζ = 23 ± 3 mV) for carboxylated MSN (MSN-) and aminated MSN (MSN+), respectively (**Fig. 2D**).

We have shown that the addition of organosilane is a limiting step in this synthesis. We have assessed different conditions for the organosilane time of addition (after TEOS) and found that a delayed co-condensation strategy succeeds in both attaching the organosilane and retaining the particles morphology, whereas an *in-situ* co-condensation consisting of adding TEOS along with organosilane before the particle formation helps create connected pods with a final hydrodynamic size ∼ 450 nm (Fig 2B, C).

### 2.2. Design of ILMs

The implementation of the lipid-silica core shell hybrid nanoparticles involves two parallel processes: fabrication of the liposomes and MSN synthesis. The optimization of each component was carried out independently.

#### 2.2.1 Refine the liposomes

We have previously shown preferential uptake of MPL-presenting silica nanoparticles^23^ and enhanced phagocytosis of MPL-bearing silicon microparticles by myeloid cells^31^. ILM formulations were previously designed and characterized to optimally integrate MPL into the lipid layer for surface presentation. In order to harness the properties of MSN for cargo loading, eighteen immunogenic lipid (IL) formulations (Table S1) were created and evaluated (Table S2) to determine optimal negatively charged and positively charged lipid compositions capable of creating colloidally stable and homogeneous PEG-free adjuvanted ILMs. The dismissing of PEG from the formulation is based on data supporting that PEG reduces uptake in immune cells (CITE) and is used in vivo as a stealth agent^32, 33^. Liposomes consisted of a mixture of up to four lipids including amphiphilic skeleton lipids DPPC or EGGSM, helper lipids as DOPC and DOPE, fluidity influencer cholesterol, and charged lipids including anionic DMPG or cationic DOTAP, DSTAP, or DODMA before incorporation of MPL at dosage up to 2%mol (∼5% wt) to endow liposomes with an immunogenic character (ILs) (Tables S1, S2).

The homogeneity of ILs was evaluated by their hydrodynamic diameter and the corresponding polydispersity indices as well as the zeta potential (Table S2). We noted the correlation of the incremental increase of the aforementioned lipids with the size and charge of the liposomes (ranged from 24 to 91 nm and PDI <0.33), except the formulation consisting of pure DPPC that showed polydisperse populations with 195 nm and PDI = 0.59, indicating that cholesterol was required in the subsequent formulations.

Next, all eighteen formulations were fused on MSN to create ILMs using ovalbumin as a model antigen, for which the cut-off size was 250 nm and PDI = 0.25 indicative of an acceptable degree of sample homogeneity. This size cut-off is justified by our envisaged intraperitoneal administration approach to promote ILM accumulation in tumor-burdened tissues. Entries in table S2 were split between ILM-(IL-fused on MSN+) and ILM+ (IL+ fused on MSN-) and were highlighted in dark red/green (No-go/Go) and light red/green for formulations that showed promise before selecting the formulations #8 fused on positively charged MSN (ILM8: 230 ± 4 nm, PDI = 0.14 ± 0.2, ζ = -28 ± 0.5 mV) and #18 fused on negatively charged MSN (ILM18: 215 ± 9 nm, PDI = 0.20 ± 0.01, ζ = 8.9 ± 0.4 mV) to pursue the study (**Fig. 2D** **and Table S2**). The ILM8 and ILM18 will be respectively denoted ILM- and ILM+ thereafter.

The hybrid particles with biomolecules and lipids were imaged by both cryoEM showing the surrounding shell indicative of presence of the lipid/biomolecule shell, and phosphotungstic acid-staining where ILM-showed a dark halo (∼5-8 nm thick) visually indicative of the lipid coating over MSN (**Fig. 2E**) comparable with our previous observations ^15, 34^. The Thermal gravimetric analysis (TGA) performed in air showed weight loss attributable to 45% lipids (**Fig. 2F**). Therefore, each 1 mg MSN would be coated by approximately 900 µg lipids, equivalent to 45 µg MPL/mg MSN. The bare MSN weight loss (∼ 10%) is attributed to the continued matrix condensation reactions with increasing temperature that evolve water and ethanol.

#### 2.2.2. Strategies to create ILM+ and ILM-with a dual adjuvant (loading and release)

Next, the selected mono-adjuvanted ILM- and ILM+ containing MPL were then optimized to load CpG in the presence of OVA as a model antigen. The combination of MPL and CpG activates both surface TLR3 and endosomal TLR 9 on targeted antigen presenting cells.^35, 36^ Considering the hierarchy of the construct in terms of type, charge and size diversity of cargo, as well as the charge of both the MSN and the liposomes, we followed various strategies depicted in **Figures 3A and 4A**, consisting of modifying the incubation order of CpG and OVA as well as that of the lipids fusion to minimize electrostatic repulsive forces and maximize loading. The loading of CpG and OVA was carried out by fluorescence reading of the dye-tagged biomolecules (CpG-FITC and OVA-TX red) in separate experiments to avoid any possible FRET-based false signal if both fluorescently tagged molecules were used simultaneously.

**Figure 3.**
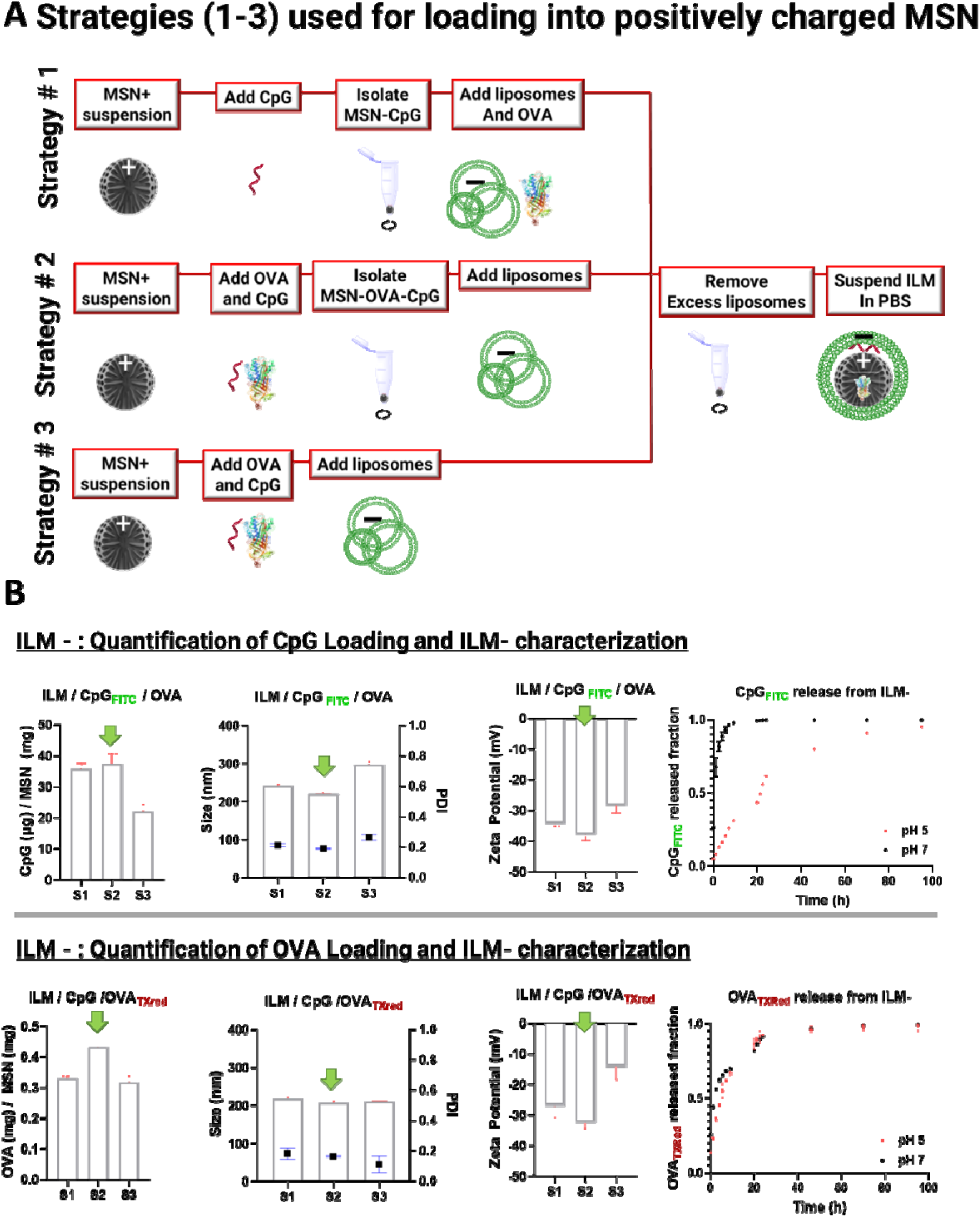
A) schematic of the different strategies (# 1,2, and 3) used to create ILM- and B) characterization of loading%, size, zeta potential of ILM- and release of CpG and OVA from dual adjuvanted ILM- made with OVA and CpG adjuvant. Green arrows indicate the selected strategy.

**Figure 4.**
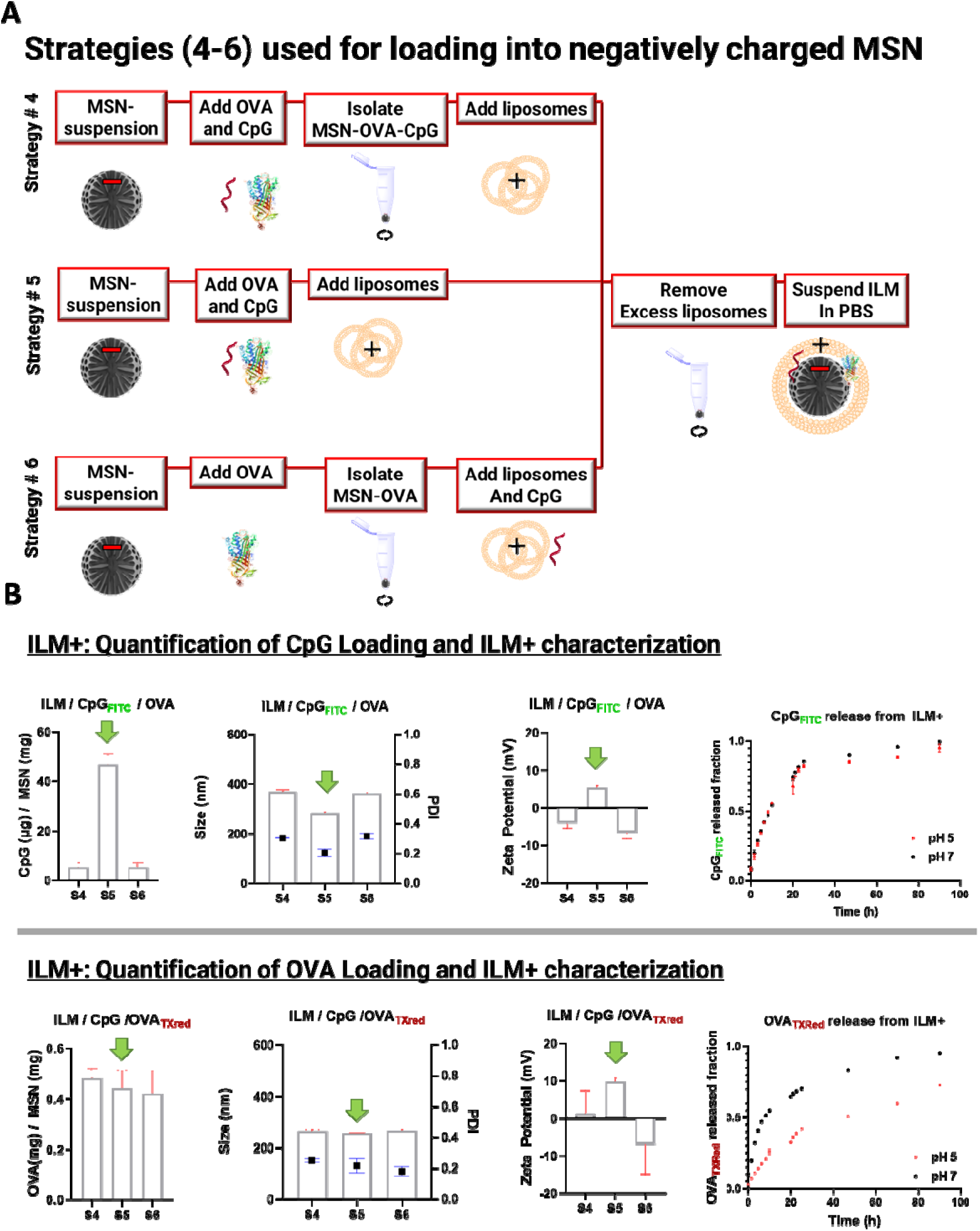
A) Different strategies (# 4,5, and 6) used to create ILM+ and B) characterization of loading %, size, zeta potential of ILM+ and release of CpG and OVA from dual adjuvanted ILM+ made with OVA l and CpG adjuvant. Green arrows indicate the selected strategy.

#### Strategies for ILM- (OVA+MPL+ CpG)

First, for ILM- (**Fig. 3**), we show that strategies 1 and 2 were comparable where negatively charged CpG was loaded in a comparable fashion into positively charged MSN independently to the loading conditions. Strategy 2 yielded slightly smaller size, and more OVA being loaded. we found that loading CpG and OVA together on positive MSN followed by isolating CpG-OVA-MSN before the addition of ILs (strategy 2, **Fig. 3A**) loaded maximum of 75% CpG (38 µg) and 90% (450 µg) OVA per mg MSN. Other strategies led to either loss of CpG retention or failure to yield a positively charged ILM (**Fig. 3B**). Strategy 3 was dropped as the simultaneous addition of negative liposomes with the biomolecules competed with the loading of CpG and yielded a two-fold decrease in the CpG loading.

#### Strategies for ILM+ (OVA+MPL+ CpG)

In the case of ILM+ exhibiting a negative MSN core and a positive lipid shell (**Fig. 4**), due to the overall positive charge of OVA in the loading conditions, we hypothesized that OVA can independently interact with CpG and drive its loading to MSN (strategy 4, **Fig. 4A**). However, CpG loading was negligible in this case likely because the positive OVA preferentially interacted with the negative silica surface instead of driving CpG onto MSN. We found that the simultaneous addition of the positively charged lipids along with negatively charged biomolecules (strategy 5, **Fig. 4A**) drive those latter onto the negatively charged surface of MSN, resulting in 80% CpG (40 µg) CpG and 90% (450) µg OVA loading (**Fig. 4B**). Although we had proven a CRISPR ribonucleoprotein driven loading via this strategy^34^, this is the first time we showcase the success of this strategy on a dual loading of both protein and nucleic acid. Strategy 6 that yielded negative nanoparticles was dropped due to the likely association of CpG with the liposomes which excess is needed to fuse on MSN, which lead to spreading the added CpG amount between fused and excess (discarded) liposomes. The final negative charge suggests that the CpG was on the outer surface of the nanoparticles rather than inside the construct.

As a control experiment, we assessed the fusion of positive ILs on the positive MSN and obtained >500 nm and PDI>0.4 indicating large and polydisperse systems beyond our cut-off threshold indicated earlier in (**Fig. S1**).

The release profiles for CpG and OVA show opposite trends for ILM+ and ILM-. The positively charged MSN in ILM-released CpG quicker at neutral pH due to stronger electrostatic interactions at acidic pH. OVA was released in a comparative manner at both pH values indicating that masking of electrostatic interactions by the ionic strength is sufficient to release the OVA model antigen from ILM-. Negatively charged MSN in ILM+ released CpG in a comparative manner at both pH values due to comparable repulsion at both pH values. Considering the OVA pI ∼ 4.5, it is understandable that at pH 7, both OVA and MSN are negatively charged promoting release more than acidic pH where OVA is almost neutral.

#### Strategies for ILM-OXA

To make OXA-loaded ILM (+/-), OXA was loaded into MSN by incubation overnight in water or DMSO solutions. The completion of the procedure followed strategies consisting of either isolating the MSN-OXA before the addition of ovalbumin, CpG and liposomes, or adding the ovalbumin and liposomes on the MSN OXA solution. Detailed procedures are shown in section 3.3.7. Oxaliplatin loading in ILM and quantification of oxaliplatin (Pt) by ICP/OES is shown in (Table S3).

### 2.3. ILM MPL-CpG and subcomponents toxicity profile with RAW264.7 macrophages and BR5 ovarian cancer cell lines

We have first assessed the safety profile of the chemo-free ILMs and their subcomponents (MSN+/-, liposomes IL+/-, non-immunogenic lipid-MSN LM+/-, and full ILM+/-) by incubating the used concentration (∼ 20 µg/mL) in the ex vivo experiments and higher concentrations (100 and 250 µg/mL) for 24 and 48 hours with 10,000 cells/well in 96 well plates and we have used Cell Titer Glo (Promega) as a bioluminescent assay for viability. BR5-akt ovarian cancer cells show low toxicity throughout the whole spectrum of the study, only ILM- and LM+ displayed cytotoxicity (viability >70%) but only at higher concentrations (100 and 250 µg/mL) and longer exposure times (48h). The RAW cells were more sensitive to these nanoparticles. Whereas ILM-does not cause any significant cell death, comparably to ILM+ at low concentration (20 µg/mL), this latter causes significant toxicity at higher concentrations (100 and 250 µg/mL) and longer exposure times (48h) closer to ∼ 50% death. After bringing the oxaliplatin-loaded ILM to the spectrum, the toxicity profile drastically changes and shows a nearly full death of RAW cells for both ILM+ and ILM-. Conversely, ovarian cancer cells were generally more resistant to OXA-ILM (except the OXA-ILM+ at 48h).

These data emphasizing the unfortunate death of myeloid cells following uptake of chemo-loaded ILM, suggesting that only drug-free ILM can support an immune response dependent on macrophages.

### 2.4. ILM MPL-CpG induced increased activation and antigen expression of DCs in-vitro

We tested the immunogenic properties of the different ILM configurations along with their sub-components ex-vivo by incubating bone marrow-derived dendritic cells (BMDC) with ILMs. Flow cytometry was used to measure the expression of costimulatory molecules (namely CD40 and CD86) indicative of cell activation, and SIINFEKL peptide presentation by major histocompatibility complex I (MHC I; H-2Kb) indicative of processing and presentation of antigen, (**Fig. 5C**). As expected, myeloid cell activation, as measured by expression of CD86 and CD40, increased the most for all ILM configurations containing MPL. The presence of CPG alone (without MPL), which is residing within the ILM core, shows negligible activation of DCs, thus MPL presented on the outer surface of ILM seems to be key in DC activation. Dual adjuvanted ILM does not show a significant increase in myeloid activation compared to the MPL-ILM. This trend was charge-independent. While dual adjuvanted ILM-similarly showed no significant compared to ILM-MPL, ILM+ did show a significant increase compared to mono-adjuvanted ILM+. It is well established that positive charge on the nanoparticle increases uptake kinetics, and the activation of macrophages as well as the antigen processing and presentation are dependent on the internalization level and kinetics.

**Figure 5.**
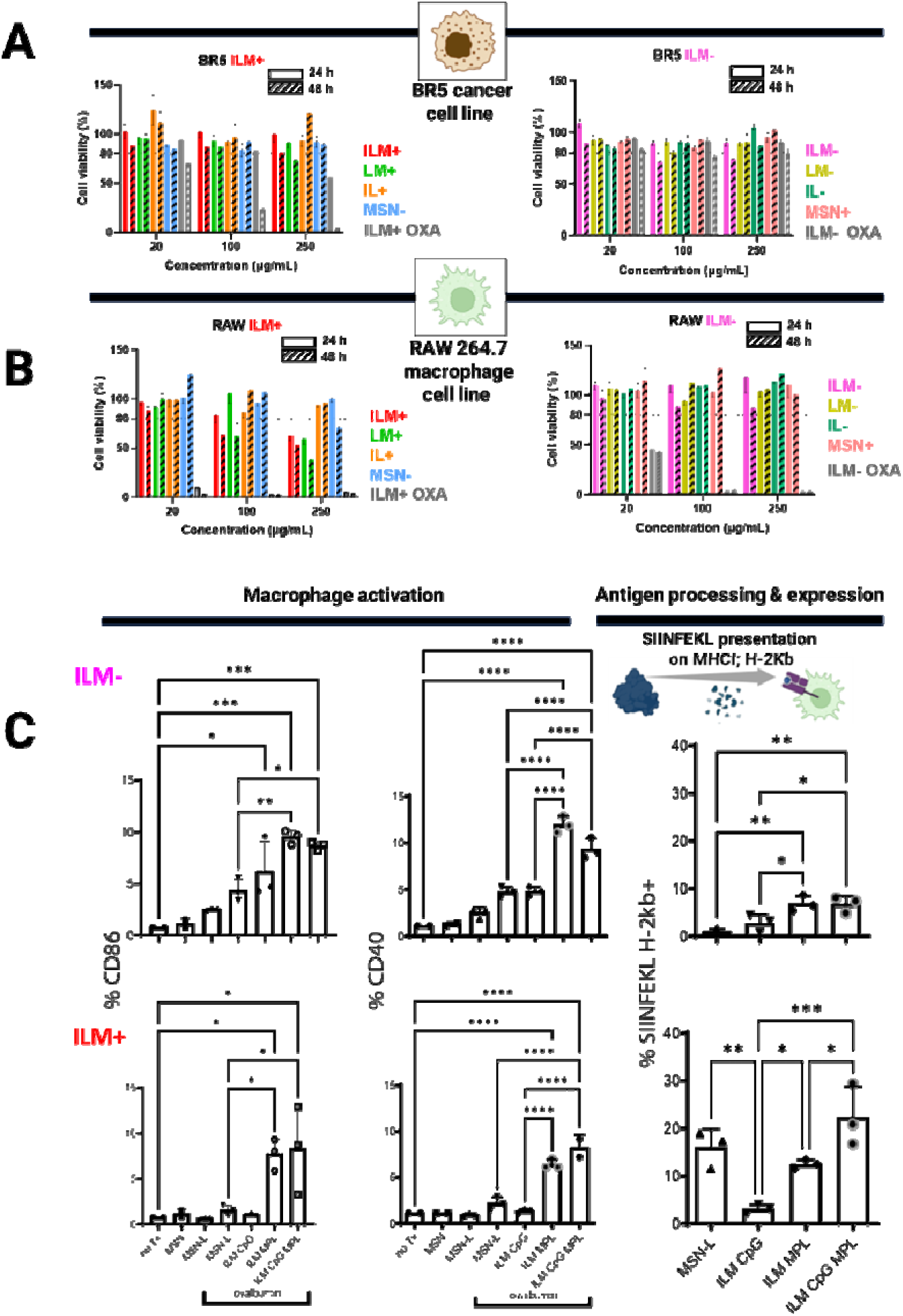
in vitro biocompatibility and activation of ILMs and subcomponents. A-B) cell viability assays for BR5 and RAW cells exposed to ILM+ and ILM- and their sub-components (MSN+/-, liposomes IL+/-, non-immunogenic lipid-MSN LM+/-, and full ILM+/-), C) flow data of expression of CD86 and CD40 activation biomarkers and SIINFEKL peptide from OVA antigen processing and presentation shown on DC

### 2.5. ILM biodistribution

In order to track mono vs dual adjuvanted ILMs *in vivo*, ovarian cancer was established in FVB mice by intraperitoneal injection of BR5-Akt-Luc cells, 250 µg fluorescent ILMs were administered intraperitoneally on Day 17, followed by whole body fluorescent imaging at 3h and 24 h later using the IVIS spectrum. While 2D imaging shows (shows ILM fluorescence specifically located within the peritoneal cavity (**Fig. 6B**), 3D imaging presents a more spatially accurate biodistribution within living animals, with similar presentation as tumor bioluminescence at 24 h (**Fig. 6A** **and Fig. S2**). Upon sacrifice and organ harvest (24 h) (**Fig. 5C**), ILMs displayed comparable distribution wherein they preferentially accumulated in omentum, fat pad and mesentery (tumor burdened tissues), with minimal signal from the liver. Several mice showed some accumulation in ovary. Building off of our recent study showing that >80% of nanoparticles accumulate in a size-independent fashion within tumor-burdened fatty tissue rather than MPS in disseminated ovarian cancer, here we show that the accumulation in those tissues is also agnostic to adjuvants and to charge (**Fig. S3**) where both mono- or dual-adjuvanted ILM+ and ILM-are also colocalized with the tumor-burdened fat pads and omentum. The presence of MPL within the outer lipid shell of both ILMs whereas CpG is encapsulated within the nanocarrier core and not on the surface, made both systems comparable to the uptake and activation of macrophages (**Fig. 4C**) hence supporting their comparable biodistribution observations.

**Figure 6.**
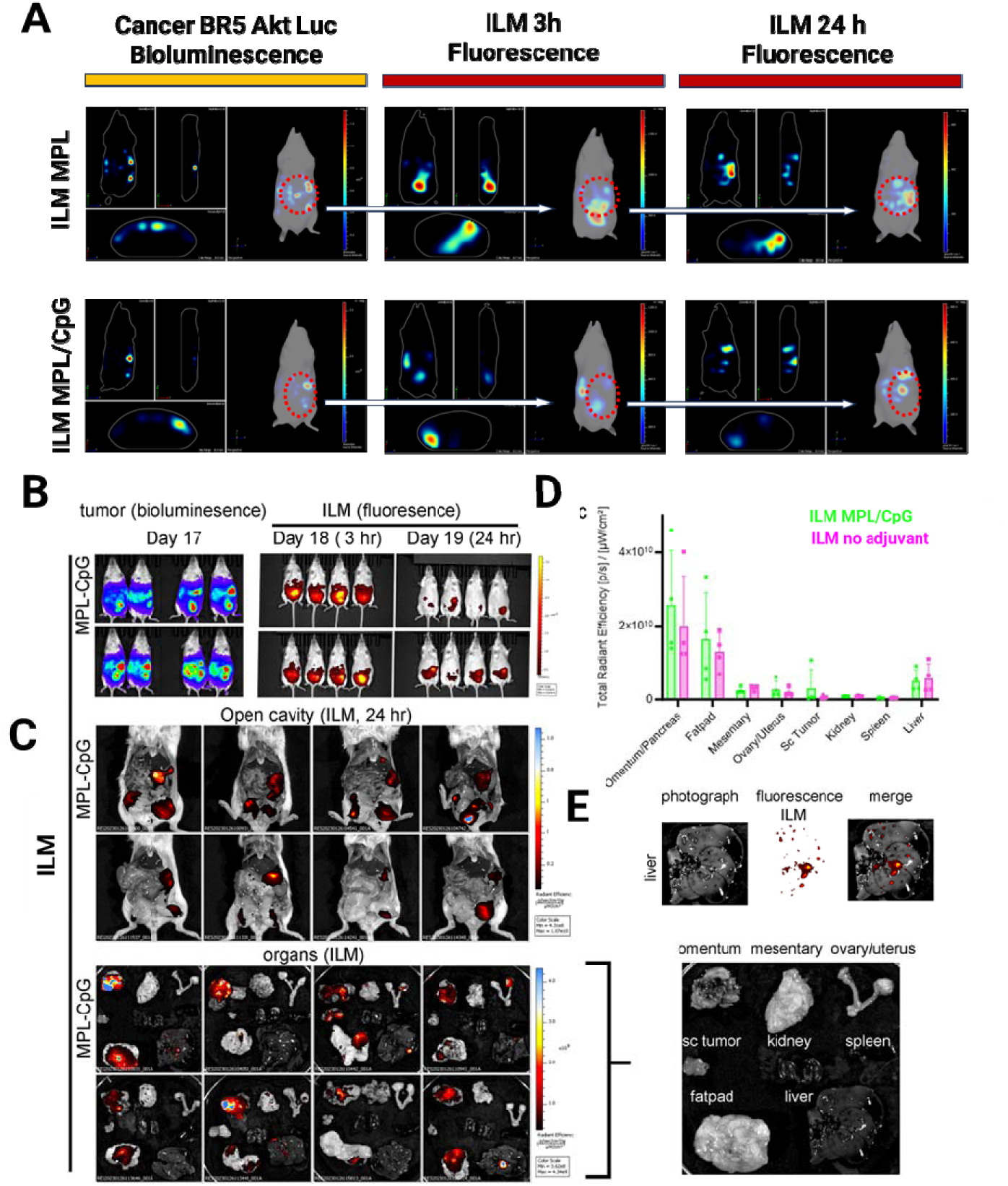
Biodistribution of ILMs in mice with BR5-Akt-Luc ovarian cancer model. A) 3D IVIS on living intact animal showing bioluminescent BR5-Akt-Luc cancer cells, and Dy633-fluorescent ILM MPL and ILM-CpG-MPL. B) 2D IVIS outlining the distribution of fluorescent ILMs in the peritoneal cavity at 3 h and 24 h after IP injection. C) imaging of sacrificed open mice with harvested organs and biodistribution of nanoparticles. Identification of organs is shown in the inset, D) total radiant efficiency showing accumulation in organs, and E) overlay of liver photograph and ILM fluorescence.

### 2.6. ILM MPL/CpG reduced tumor burden and enhanced survival

Upon cancer challenge, mice were injected on day 4 and day 11 with 200 µg ILM. To track tumor burden evolution, mice were administered luciferin solution 10 min before IVIS imaging at days 4,7,12, and 17 (**Fig. 7A**). We measured survival and tumor burden for mice receiving treatments of lipid-coated MSN (no adjuvants), ILM-MPL, and ILM-CpG-MPL, or control PBS (**Fig.7B-D**). Both ILM-MPL and ILM-CpG-MPL caused a significant reduction in tumor burden.

**Figure 7.**
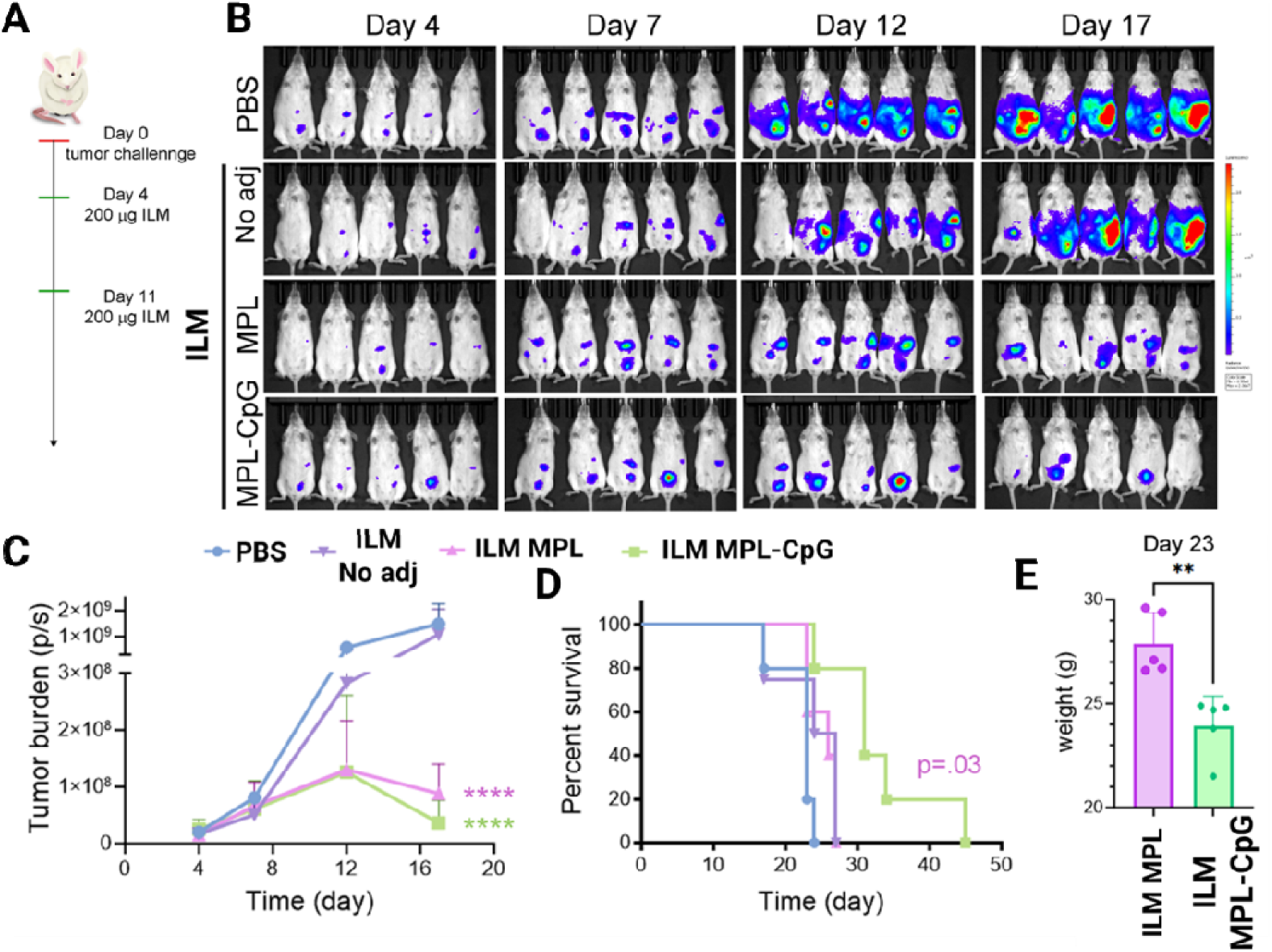
in vivo efficacy of ILM in FVB mice. A) Timeline with cancer challenge, ILM injections (treatment) and luciferase injection (IVIS), B) bioluminescent images from BR5-Akt-Luc with their C) tumor burden (photon counts) based on the bioluminescent images, D) a Kaplan-Meier survival curve are shown [Log-rank (Mantel-Cox) test] and E) mice weight on day 23 comparing ILM MPL and ILM MPL-CpG.

While ILM-MPL showed only a modest increase in survival comparable to control carrier and PBS (∼ 30 days), ILM-CpG-MPL administration led to more robust (p=0.03) survival (**Fig. 7D**) with up to 45 days (time we ended our study).

The preferential ILM colocalization with CD11b+ cells (**Fig. S4**), consistent with our previous observations^22, 23^, supports that the IP administered nanoparticles uptake within the myeloid cells is agnostic to particle features. The use of oxaliplatin within ILM was also optimized to maintain stability and loading of model antigen and adjuvants, with the preferential accumulation of ILMs in myeloid cells rather than cancer cells explaining why the simultaneous use of chemotherapy in this case is not an appropriate approach (**Fig. S5**). The possibility of in vivo nanoparticle transfer from macrophages to cancer cells via tunneling nanotubes did not appear to occur to a degree that could support selective death of cancer cells ^37, 38^.

### 2.7. Development of a Semi-mechanistic Model for NP-Driven Tumor Immunotherapy

To describe tumor regression in response to immunogenic NP therapy, we developed a semi-mechanistic model that couples tumor growth dynamics with innate immune activation using a minimal system of ordinary differential equations (ODEs; see Methods). The model tracks two variables over time: tumor burden vCiJ, which grows logistically in the absence of treatment, and a generalized immune effector population MCiJ, representing anti-tumor innate immune activity (e.g., macrophages, NK cells). Rather than explicitly modeling the full cascade of immune cell recruitment, cytokine signaling, and T cell priming, we abstract this complexity into a single effector variable. This simplification is justified by the short experimental timescale (2–3 weeks) and the lack of evidence for adaptive immune engagement during this window.

Thus, MCiJ serves as a compact but interpretable proxy for innate immune function. NP-induced immune stimulation is modeled through an external forcing function NCiJ, which introduces exponentially decaying pulses following each NP dose. These pulses mimic transient immunogenic activation and are scaled by the parameter *k_act_*, which governs the strength of stimulation per dose. In this framework, *k_act_* also captures enhancements due to tumor antigen targeting, such as the delivery of tumor antigen. Increases in *k_act_* are therefore used to simulate improvements in NP immunogenicity without explicitly modeling antigen presentation or T cell engagement. The decay of immune activation is governed by a fixed parameter λ, corresponding to an assumed immune half-life of 3 days. This value reflects the transient nature of innate immune activation and is not intended to represent NP pharmacokinetics, which is not explicitly modeled. Instead, we assume that each NP dose triggers an immediate immunogenic signal, justified by the faster kinetics of immune effector activation compared to NP biodistribution.

Finally, the model is formulated as a deterministic ODE system, without stochasticity, spatial heterogeneity, or inter-subject variability. While this limits our ability to capture mouse-to-mouse variation or local immune gradients, it enables robust fitting to population-level data and facilitates interpretable simulation of design scenarios.

### Model Calibration Reveals Immune Parameter Differences and Characterizes Treatment Response

We calibrated the semi-mechanistic model to tumor burden data from mice treated with PBS, L-MSN, L-MSN-MPL, or L-MSN-MPL-CpG NPs (**Fig. 8A**). To maintain parsimony and improve parameter identifiability, the tumor growth rate γ and carrying capacity V̂ were fixed across all treatment groups based on fits to the PBS cohort (control). This allowed us to isolate and estimate only the immune-related parameters, i.e., immune killing rate (*k_M_*), immune activation rate (*k_act_*), and immune decay rate (*d_M_*), which were expected to vary with NP formulation.

**Figure 8.**
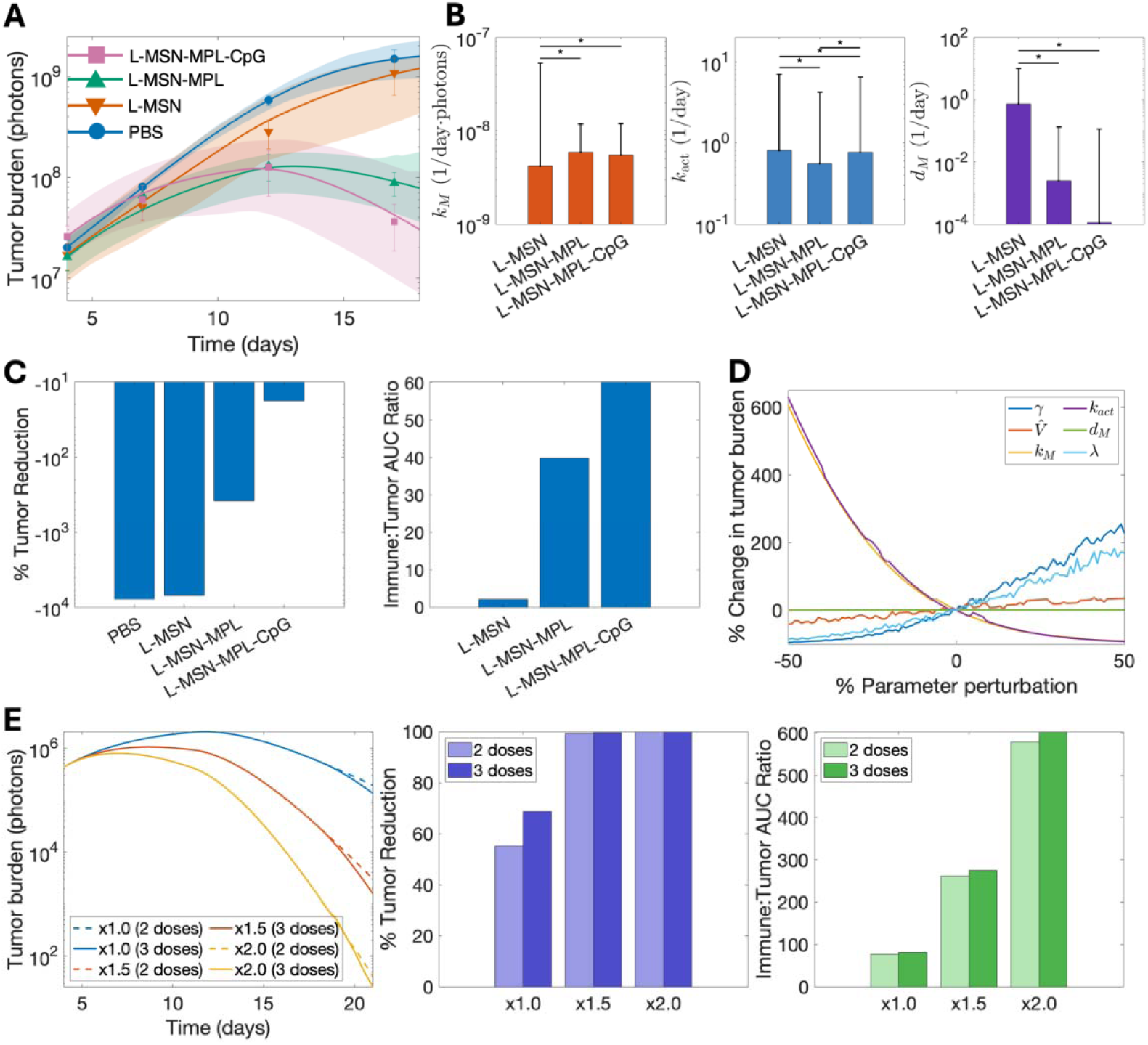
Modeling-based analysis of tumor response to immunogenic NP therapy reveals mechanistic drivers and design strategies. A) A mechanistic model was calibrated to longitudinal tumor burden data from mice treated with PBS, L-MSN, L-MSN-MPL, or L-MSN-MPL-CpG NPs. The model accurately captured the dynamics across treatment groups, with shaded bands denoting 95% confidence intervals from bootstrap fitting. B) Estimated immune-related parameters across groups: killing rate (*k_M_*), immune activation rate (*k_act_*), and immune decay rate (*d_M_*). Bars represent best-fit values with 95% confidence intervals. Asterisks indicate statistical significance from uncorrected pairwise Wilcoxon tests (*p<0.05*). C) Treatment efficacy metrics derived from the model fits. Left: Percent tumor reduction on day 18 relative to initial burden, highlighting improved outcomes with adjuvanted NPs. Right: Immune-to-tumor area-under-the-curve (AUC) ratio, quantifying immune dominance over tumor expansion. Immune AUC is not computed for PBS due to lack of immune module. D) Local sensitivity analysis showing the percent change in tumor burden on day 18 in response to ±50% perturbation in each model parameter. E) Simulated enhancement of immune activation (*k_act_*) and addition of a third NP dose on day 18. Left: Tumor burden dynamics under varying *k_act_* values and dosing schedules. Middle and right: Resulting tumor reduction and immune:tumor AUC ratio across scenarios demonstrate benefit of stronger and sustained immune stimulation.

Final parameter values and 95% confidence intervals are provided in Table S4. The model accurately captured distinct tumor trajectories across formulations, with fits tightly constrained by bootstrap-based confidence intervals. PBS-treated animals showed logistic tumor growth, while immune-activating NPs induced varying degrees of regression, with the L-MSN-MPL-CpG group exhibiting the strongest and most durable response (**Fig. 8A**).

Fitted immune parameters revealed formulation-specific differences in underlying biology **(Fig. 8B**). Compared to the L-MSN control, both adjuvanted formulations exhibited modest but significant increases in innate immune killing rate **k_M_**, though no statistical difference was observed between L-MSN-MPL and L-MSN-MPL-CpG. In contrast, the immune activation rate *k_act_* showed significant pairwise differences across all three groups, highlighting its sensitivity to both MPL and CpG inclusion. The immune decay rate d_M_ was substantially reduced in adjuvant-treated groups, consistent with more durable immune stimulation. These parameter differences provide a mechanistic basis for the observed tumor regression patterns and motivate further analysis of various model parameters as drivers of therapeutic efficacy.

Furthermore, to quantify therapeutic response, we computed two model-derived metrics: percent tumor reduction at day 18, and the immune to tumor area-under-the-curve (AUC) ratio (**Fig. 8C**). Both metrics showed a clear trend: immune potency and tumor control increased with immunostimulatory content, with MPL+CpG outperforming MPL and L-MSN. These outcomes, directly derived from the calibrated model, provide a quantitative and mechanistic basis for comparing NP formulations for their anti-tumor efficacy.

### Sensitivity Analysis

To assess the influence of individual model parameters on therapeutic outcomes, we computed the percentage change in tumor burden at day 18 in response to ±50% perturbations in each parameter, relative to baseline predictions (**Fig. 8D**). This output metric reflects the sensitivity of tumor response to small local changes in model inputs, enabling identification of the most influential parameters. Our perturbation-based local sensitivity analysis revealed that tumor burden is most sensitive to changes in the immune activation rate constant *k_act_* and the immune killing rate **k_M_**. Specifically, increasing *k_act_* resulted in a sharp nonlinear decrease in tumor burden, highlighting immune stimulation as a key lever for therapeutic efficacy. In contrast, tumor growth rate v and the exponential decay rate λ associated with immune activation pulses

showed positive correlations with tumor burden, suggesting that slower tumor proliferation and slower decay of immune activation following NP dosing led to improved tumor control by sustaining post-dose immune signaling. Notably, the decay rate of immune effectors d_M_ and the tumor carrying capacity V̂ had minimal influence on model output within the perturbation range, implying these parameters are less critical for short-term tumor response under the current therapeutic scenario. Among all parameters, *k_act_* is of particular interest due to its experimental tunability. Unlike **k_M_**, which reflects intrinsic immune cell cytotoxicity and is not easily modifiable, *k_act_* can be enhanced through NP design, such as incorporating potent adjuvants (e.g., CpG) or functionalizing particles with tumor antigens to promote antigen-specific immune

activation. These insights motivated our subsequent analysis (**Fig. 8E**) exploring how modulating *k_act_* and NP dosing frequency can be leveraged to improve therapeutic outcomes.

### Modulating Immune Activation and Dosing Frequency Enhances Tumor Control

Building on the sensitivity analysis identifying *k_act_* as a key determinant of therapeutic efficacy, we simulated tumor response under hypothetical modifications to NP design and dosing schedule (**Fig. 8E**). Specifically, we increased *k_act_* by 1.5× and 2.0× relative to baseline to reflect enhanced immune activation, such as might be achieved by tumor antigen targeting or adjuvant optimization, and evaluated outcomes under both carried out 2-dose and theoretical 3-dose regimens.

The resulting tumor burden trajectories (left panel) demonstrate a dose- and potency-dependent improvement in tumor control. While a modest increase in *k_act_* (1.5×) reduced tumor size substantially with two doses, a third dose, administered on day 18, one week after the second dose, consistent with the experimental protocol, further accelerated regression. At 2.0× *k_act_*, tumor elimination was nearly complete by day 21, with diminishing marginal benefit from the third dose. These findings underscore the synergistic potential of combining stronger immune stimulation with repeated administration.

This trend was further reflected in the calculated efficiency metrics (middle and right panels). Percent tumor reduction increased sharply with higher *k_act_*, saturating near 100% for both dosing scenarios at 1.5× and above. Similarly, the immune to tumor AUC ratio, a proxy for treatment efficiency, rose monotonically with increasing *k_act_* and was further amplified by a third dose. These simulations reinforce that enhancing the magnitude and duration of immune

activation is a powerful strategy for improving outcomes, particularly in regimens constrained to short treatment windows.

Importantly, these results also emphasize that additional dosing alone is insufficient if immune activation is weak, thereby highlighting the need for immunogenic NP formulations capable of robust immune engagement. Together, these findings support a rational design framework in which mechanistic modeling guides the optimization of NP potency and scheduling for maximal therapeutic impact.

## Conclusion

In summary, we have assembled a nanoparticle platform that successfully concentrates multiple active complementary biological components with different natures and sizes (protein, oligonucleotide adjuvant, lipid adjuvant and small molecule drug). The platform consists of a hybrid silica-lipid nanoparticle which refined features and optimized strategies of loading allowed its successful application to ovarian cancer immunotherapy. The rational design of the platform has considered the major factors of electrostatic interactions between the silica core, the cargo, and the lipid shell, the lipid composition and the co-loading of immune adjuvants coactivating multiple TLRs. The lipid shell was optimized to contain the lipid-based adjuvant MPL while keeping the particles homogeneity and colloidal stability allowing for an optimal strategy to incorporate CpG deoxyoligonucleotide (TLR3/9 adjuvant) in a final construct which final charge is tunable in a straightforward fashion. The immunogenic lipid-MSN (ILMs) successfully activate myeloid cells as demonstrated by an increase in CD40 and CD86 expression in an MPL- dependent fashion. Antigen processing and presentation was optimal using the dual adjuvanted ILM+, likely due to enhanced internalization by myeloid cells. As such, the addition of chemotherapy in this system was counterproductive. ILMs were predominately located in tumor-burdened tissues in an adjuvant- and charge-independent fashion. The dual adjuvanted ILM- caused a significantly higher probability of survival compared to mono-adjuvanted sample. Amongst ILM, chemo-free dual adjuvanted nanoparticles with a final positive charge appears to be an optimal system to induce an efficient immune response against ovarian cancer, and this was supported by a semi-mechanistic model which quantified treatment efficacy and identified key parameters governing tumor response hence guiding the future design of nanoparticle-driven tumor immunotherapy

## Conflict of interest

Authors have no conflict of interest to declare.

## Acknowledgement

The authors were grateful for assistance and use of the University of New Mexico Comprehensive Cancer Center Animal Models, Fluorescence Microscopy, Flow Cytometry and Histology Shared Resources, supported by NIH grant NCI P30 CA118100 (PI Sanchez). This work was also supported by the Oxnard Foundation (PI Serda, R) and by the AIM center core funded by NIH grant P20GM121176. Research reported in this publication was also supported by National Institute of Biomedical Imaging and Bioengineering of the National Institutes of Health under award number R01EB035545 (MPIs Dogra, Noureddine). The content is solely the responsibility of the authors and does not necessarily represent the official views of the National Institutes of Health. We would like to thank the Center for Metals and Biology and Medicine pilot funding from P20GM130422-03. We gratefully acknowledge use of the UNMCCC Animal Models, Fluorescence Microscopy, Flow Cytometry, and HTR Shared Resources as well as the NIH P30CA118100 grant that supports the UNMCCC and these shared resources.

This work was performed, in part, at the Center for Integrated Nanotechnologies, an Office of Science User Facility operated for the U.S. Department of Energy (DOE) Office of Science. Los Alamos National Laboratory, an affirmative action equal opportunity employer, is managed by Triad National Security, LLC for the U.S. Department of Energy’s NNSA, under contract 89233218CNA000001. Sandia National Laboratories is a multimission laboratory managed and operated by National Technology and Engineering Solutions of Sandia, LLC, a wholly owned subsidiary of Honeywell International, Inc., for the U.S. Department of Energy’s National Nuclear Security Administration under contract DE-NA0003525.

## 3. Experimental Details

### 3.1. Equipment

TEM images were acquired on a JEOL 2010 (Tokyo, Japan) instrument equipped with a Gatan Orius digital camera system (Warrendale, PA) under a 200 kV voltage. Hydrodynamic size and zeta potential data were acquired on a Malvern Zetasizer Nano-ZS equipped with a He–Ne laser (633 nm) and non-invasive backscatter optics. All samples for DLS measurements were suspended in distilled water or ethanol at a 1 mg mL^−1^ concentration. Samples were washed three times through centrifugation prior to measurements. Measurements were acquired at 25 °C. DLS measurements for each sample were obtained in triplicate and then the Z-average diameter (by Intensity) was used for all reported hydrodynamic size values. The zeta potential for all the samples was measured in distilled water in triplicates according to Smoluchowski theory. All reported values correspond to the average of three independent measurements. TGA was carried out by TA instruments, samples were placed in alumina crucible and burned up to 650 °C under 25 mL/min air flow. Plate reader was used to evaluate loading of biomolecules via fluorescence and was BioTek Synergy H1 using disposable 96-well opaque microplates. For cryo-TEM sample preparation the solution must be vitrified in place in amorphous ice using a commercial or custom in-house vitrification instrument. The samples are plunge frozen in liquid ethane and stored in liquid LN_2_ before imaging, with the exact procedure depending on the specific setup for vitrification. For the images in this paper a Thermo Fisher MKIV Vitrobot was used, with the samples imaged on a Talos L120C cryo-TEM using a Gatan 626 holder. Thermo Fisher Attune NxT flow cytometer was used, and analyses used SpectroFlo Flow Cytometry Software v3.0 (Cytex Biosciences, Fremont, CA, USA) and FlowJo (10.6) (Becton, Dickinson and Company; Franklin Lakes, NJ, USA).

### 3.2. Materials

#### Antibodies

CD11c PE (N418), CD86 eFluor 450 (GL1), and SIINFEKL H-2kb APC (eBio25-D1.16) were purchased from eBioscience/Thermo Fisher Scientific. CD40 FITC (3/23) was purchased from BD Biosciences.

#### Cells Lines and Mouse Models

The BRCA1-deficient BR5-Akt cell line, generated on an FVB background, was a kind gift from Dr Sandra Orsulic (Cedars-Sinai). To monitor tumor burden using a bioluminescent tag, the cell line was lentivirus transduced to constitutively express firefly2 luciferase (Br5-*akt*-Luc). Cell lines were cultured in DMEM containing 10% FBS and 100 units per 100 µg penicillin/streptomycin at 37 °C and 5% CO2. Trypsin-EDTA was used to collect cells. To prepare BMDC, bone marrow was collected from the femurs of female murine C57BL/6 or FVB mice using a 27 g needle and syringe to flush the marrow from the bone. RBC were lysed with BD lysis buffer as described by the vendor. Cells were cultured in 6-well plates (3 ml per well) for 8–10 days in RPMI 1640 medium supplemented with 10% FBS, 100 mM β- mercaptoethanol, penicillin/streptomycin and 10 ng/mL recombinant murine GM-CSF. Half of the media was replaced every 2–3 days with fresh media and cytokines. Mice were purchased from Charles River or Jackson Laboratories and housed in a specific pathogen-free facility. All animal protocols were approved by the Institutional Animal Care and Use Committee (IACUC) at the University of New Mexico (Albuquerque, NM, USA). Mice were killed when moribund or when weight reached 30 g due to ascites accumulation. Mice were monitored and weighed every 2-3 d.

### 3.3. Methods

#### 3.3.1. Synthesis of monodisperse MSNs with ∼7 nm dendritic pores: carboxylated (MSN-) or primary amine-terminated (MSN+)

The synthesis of the MSNs was adapted from previously reported^29, 39^. In a 100 mL round bottom flask, triethanolamine (0.18 g) was added along with CTAC (24 mL, pH = 6) and water (36 mL). The pH of the solution was adjusted to 8.5 with sodium hydroxide and the mixture was heated to 50 °C and stirred (600 rpm) for 1 h. The stirring rate was adjusted to 350 rpm and a 20 mL solution of TEOS in cyclohexane (10% v/v) was slowly added to form a biphase system. For carboxylated MSN, triethoxypropylsuccinic anhydride (5% mol compared to TEOS) and for functionalized MSN, APTES (110 µL, for MSN-NH2) in ethanol (200 µL) were added next day (t= 16 h) in the bottom aqueous phase (containing silica nanoparticles) and kept reacting for 4 h. The upper organic phase was then removed, and the nanoparticles suspension was centrifuged. The isolated pellet is suspended in ethanol and centrifuged twice. The surfactant removal was achieved by successive washing steps in HCl (1% ethanol, twice); each step included 15 min sonication and centrifugation. All centrifugation cycles were done at 50K rcf for 20 min at 18 °C. Lastly, the template-free MSN were washed twice in ethanol and stored as a suspension in ethanol. The suspensions are stable for at least one year.

#### 3.3.2. Conjugation of Fluorescent labels on MSN

DyLight 488-NHS ester, or DyLight 633-NHS ester were used to fluorescently label MSNs. A solution of dye in DMF (1 mg/mL, 250 µL) stored at -20 °C was added to a suspension of MSN- NH2 (10 mg, 2.5 mg/mL) and reacted for 18-24 h at room temperature in the dark. To obtain negative MSN, the mixture was centrifuged and resuspended in succinic anhydride solution in DMF (100 mg, 25 mg/mL) and reacted for 24 h at room temperature in the dark (in order to turn the remaining amines into amides). To keep the particles positive, this step is skipped. Next, the mixture was centrifuged, and the isolated dyed pellet was washed in DMF (once) then in pure ethanol (twice). An aliquot was washed in water twice to confirm the final charge of the fluorescently labeled MSN.

#### 3.3.3. Preparation and assessment of immunogenic liposomes

We have prepared a library of 18 lipid formulations (Table S1) containing 0-2 %mol MPL. The components used in this library are cholesterol, 1,2-dipalmitoyl-sn-glycero-3-phosphocholine (DPPC), Egg Sphingomyelin (Egg SM), 1,2-dioleoyl-sn-glycero-3-phosphoethanolamine (DOPE), 1,2-dioleoyl-sn-glycero-3-phosphocholine (DOPC), 1,2-dimyristoyl-sn-glycero-3- phospho-(1’-rac-glycerol) (sodium salt) (DMPG), 1,2-dioleoyl-3-trimethylammonium-propane (chloride salt) (DOTAP), 1,2-stearoyl-3-trimethylammonium-propane (chloride salt) (DSTAP), 1,2-dioleyloxy-3-dimethylaminopropane (DODMA) (Avanti Polar Lipids) and MPL lipid A was from Millipore Sigma.

Individual lipids in chloroform (1, 10 or 25 mg/mL) are stored under argon atmosphere at -25°C. the 18 PEG-free lipid formulations were prepared by mixing the corresponding constituting lipids in a glass vial (in a glovebox) with total lipid mass per vial ranging from 5 to 20 mg depending on the experiment scale. The chloroform was removed from the lipid mixture under reduced pressure (rotator evaporator, 5 minutes). The dry lipid film was then hydrated using 1xPBS to 5 mg/mL and sonicated for at least 15 minutes.

#### 3.3.4. MPL incorporation

MPL adjuvant was incorporated within the liposomes at 2% mol, equivalent to 5% weight. The TGA analysis indicates that for each 1 mg MSN, there is around 900 µg lipids equivalent to 45 µg MPL.

#### 3.3.5. Loading of Ovalbumin and CpG, and ILM assembly

The loading of OVA and CpG adjuvant was carried out by assessing different strategies to maximize the loading of both biomolecules to obtain both positively and negatively charged ILMs. The strategies are shown in Figures 3A **and 4A** and consist basically of modifying the order of biomolecules addition and particle isolation. In summary, MSN suspension (200 µg, 1 mg/mL) was incubated with CpG (10 µg, 0.1 µg/µL) alone (strategy 1) or with OVA (100 µg, 1 mg/mL) (strategy 2 and 3) for 15 minutes before centrifugation for 10 minutes at 20 Krcf. Liposomes (5 mg/mL, 1 mg) (with OVA, strategy 1), or alone (strategy 2 and 3) is added to the loaded MSN suspension and the mixture is briefly sonicated and centrifuged (10 min, 20 Krcf) to remove excess liposomes, before being suspended in PBS at 1 mg per mL.

To quantify OVA and CpG loading, the fluorescently labeled version of each of the biomolecules is used, and the cumulated supernatants as well as the MSN suspension are saved for loading assessment by fluorescence measurement.

#### 3.3.6. OVA and CpG loading and release quantification

OVA TX red (used with non-fluorescent CpG) CpG FITC (used with non-fluorescent OVA) Loading of OVA and CpG was assessed by emission fluorescence using plate reader. Fluorescently tagged molecules, namely OVA-Texas Red (Ovalbumin, Texas Red™ Conjugate, ThermoFisher Scientific) and CpG-FITC (ODN 1826 FITC, invivogen) were used and the loading was assessed as follows:

-Texas Red-BSA - Excitation: 530nm, Emission Start/Stop: 560-700nm
-FITC CpG - Excitation: 400 nm, Emission Start/Stop: 430-700nm

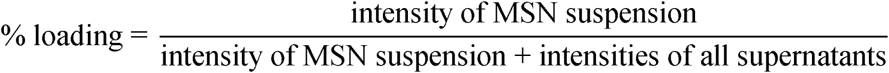

##### Release Studies

The release of CpG and OVA from ILM- and ILM+ were carried out by incubating ILMs loaded with both molecules but alternating the fluorescent cargo between CpG-FITC (invivogen) and OVA-TXred (ThermoFisher Scientific). Upon incubation at 0.5 mg/mL ILM in PBS (neutral pH) or MES buffer (pH 4.7) at 37 °C under rotating conditions (60 rpm), at different times (t) the ILM’s suspension fluorescence was recorded, then the ILM was isolated by 15K rcf centrifugation (10 min) and the fluorescence of the supernatant was recorded using exactly the same conditions. The release at each time (t) was calculated as a ratio of fluorescence from supernatant over the suspension.

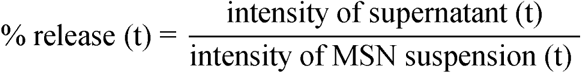

#### 3.3.7. Oxaliplatin loading in ILM

OXA was loaded into MSN by incubation overnight in water or DMSO solutions. The strategies 1 and 2 consist of:

1. isolating the MSN-OXA before the addition of ovalbumin and liposomes, or
2. adding the ovalbumin and liposomes on the MSN OXA solution

Liposomes were prepared by sonication in PBS (1 mL containing 1 mg OXA) following these strategies:

1 – MSN- : add OVA and CpG together, followed by addition of L+ (without any middle stage separation).

2- MSN+ : add OVA and CpG, spin down, add L+ on the suspension.

Then, ICP-OES was used to measure oxaliplatin loading by assessing Pt concentration nanoparticles (Table S3). ILM loaded with Pt were exposed to aqua regia (1:3 mixture of ultrapure HNO3 and HCl) with a Digi prep MS SCP Science block digester at 95 °C for 4 h. The digested samples were diluted and passed through 0.45 μm filter. The concentration of silicon was then measured using a Perkin Elmer Optima 5300DV ICP-OES, with a detection limit of <0.5 mg l^−1^. ICP-OES was calibrated with a five-point calibration curve. QA/QC measurements were also obtained to ensure quality results.

***Quantification of Pt in ILM-oxaliplatin:*** OXA@ILM- and OXA@ILM+ obtained upon loading were acid digested for ICP-MS to quantify the Pt % in function of the loading conditions. ILM- exhibited the highest loading of OXA with up to 5.2% (=2.6% Pt found by ICP-MS) without alteration of the OVA and CpG loading.

#### 3.3.8. In vitro DC functional studies

Bone marrow-derived DCs were seeded in 12-well plates at a density of 1 × 10^5^ cells per well (1 mL). After 24 h, the media was removed and replaced with 2 ml of fresh complete media supplemented with 25 µg MSN, lipid-coated MSN, ILM-CpG, ILM-MPL, or ILM-CpG-MPL and incubated for 24 h. DCs were collected using 3 mM EDTA. The suspended cells were centrifuged, washed with PBS containing 1% BSA and labelled with fluorescent antibodies specific to CD11c, CD86, CD40, and SIINFEKL H-2kb. Cells were analyzed using the Thermo Fisher Attune NxT flow cytometer.

#### 3.3.9. Proliferation assays

ILM and their subcomponents at different concentrations were assessed for cell compatibility using the CellTiter-Glo 2.0 assay. Briefly, cells BR5 and RAW were seeded at a density of 100,000 cells ml^−1^ in culture media in opaque dark 96-well plates. After 24 h and 48h, CellTiter-Glo 2.0 reagent (100 µL) was added to each well, and following a 10 min incubation, luminescence was determined using a BioTek Synergy H1 microplate reader. Percent cell viability was calculated relative to control, non-treated cells.

#### 3.3.10. Imaging Tumor Burden and Biodistribution

For in vivo monitoring of tumor burden, mice with BR5-Akt-Luc tumors were administered 150 mg luciferin kg^−1^ by IP injection. Mice were then anesthetized using 2.5% isoflurane, and after 10 minutes, 2D and 3D bioluminescence images were acquired using the Xenogen IVIS Spectrum animal imager (Perkin Elmer). ROI total flux (photons/sec) measurements were acquired using Living Image Software (Perkin Elmer).

Eighteen or twenty-one days post IP BR5-Akt tumor cell injection, mice were administered 200 µg DyLight633 far red fluorescent ILMs by IP injection. Mice were imaged 24 hours after nanoparticles injection for bioluminescence (tumor) and fluorescence (MSN) using the 2D and 3D IVIS Spectrum or at different times as indicated (days 4,7,12, and 17). Peritoneal organs were harvested and imaged. ROI measurements were acquired using Living Image Software.

#### 3.3.11. Single-Adjuvant and Dual-Adjuvant Efficacy Studies

Female FVB mice (6-8 weeks old) in groups of 5 were injected intraperitoneally (IP) with 2 x 10^5^ BR5-Akt-Luc BRCA1-deficient serous epithelial ovarian cancer cells in 200 µL PBS. Four days post tumor challenge (Day 4), mice were administered with IP injection of one of four treatments: 200 µL PBS alone, adjuvant-free ILM at 200 µg/200 µL PBS, MPL-ILM at 200 µg/200 µL PBS, or MPL-CpG-ILM at 200 µg/µL PBS. This process was repeated on the eleventh day (Day 11) post tumor challenge. On the fourth, seventh, twelfth-, and seventeenth- days post tumor challenge (Days 4, 7, 12, 17), the mice were injected IP with 150 mg/kg D-luciferin potassium salt for bioluminescent imaging using the Xenogen IVIS® Spectrum (Perkin Elmer, Waltham, MA, USA). Mice were then anesthetized using 2.5% isoflurane and ten minutes after injection 2D/3D bioluminescent images were taken of the model groups. Regions of interest (ROI) were created around each mouse, and total photon counts per second (p/s) were determined and compared across groups. Mouse weight was tracked, along with abdominal width and mouse appearance. Critical endpoints for sacrifice included: 30 g with ascites present, and ruffled fur and unhealthy appearance. ^26^

#### 3.3.12. Mathematical Modeling of Nanoparticle-Driven Tumor Immunotherapy Response

To quantitatively capture the observed tumor response to immunogenic NP therapy, we adapted and extended a mechanistic modeling framework previously developed by our group for evaluating tumor dynamics in therapeutic settings^40–45^.The model accounts for canonical tumor growth kinetics and extended to incorporate NP-induced activation of effector immune cells, primarily innate effectors such as macrophages or NK cells, assumed to be responsible for tumor cytotoxicity in this context.

The model comprises two state variables: tumor burden (vCiJ) and an abstracted immune effector variable (MCiJ), representing activated innate immune response. Tumor growth in the absence of treatment follows logistic growth, and immune-mediated tumor killing is represented as a bilinear interaction, which assumes tumor death is proportional to both tumor and effector cell abundance. Thus, we get:

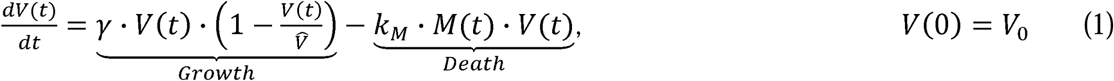

where γ is the intrinsic tumor growth rate, V̂ is the tumor carrying capacity, and **k_M_** is the cytotoxic efficiency of the immune response.

Effector cell dynamics are governed by NP-induced immune cell activation and natural decay:

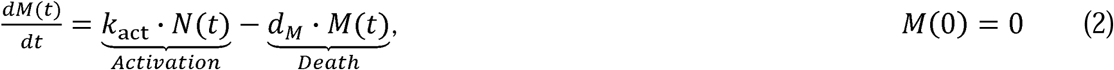

Here, *k_act_* quantifies the strength of NP-induced immune activation, and d_M_ denotes the degradation or exhaustion rate of effector cells. The forcing function NCiJ captures the time-dependent immunostimulatory input resulting from administered NP doses. Each NP injection is modeled as generating an exponentially decaying immune activation pulse. Specifically, we define:

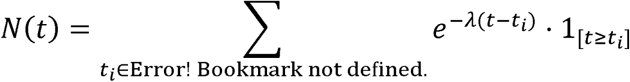

where i_i_ denotes NP injection time points (e.g., days 4 and 11), and 1_[*t*≥*t*_i__] is the indicator function, equal to 1 when *t*≥*t*_i_ and 0 otherwise. λ - log(2)/*t*_1/2_ reflects the decay rate corresponding to an immune activation half-life of 3 days. This half-life was chosen based on the time window over which innate immune signaling and tumor regression were observed in vivo.

This formulation ensures that each NP dose triggers an exponentially decaying immune activation pulse starting at its respective injection time. The additive structure of NCiJ implies that immune stimulation is both independent and non-saturating across doses. That is, each dose is assumed to elicit a consistent level of immune activation, regardless of the timing or magnitude of previous doses. This simplification avoids the complexity of modeling dose-to-dose interactions such as tolerance, synergy, or saturation. It is justified by the use of submaximal NP doses in our study, where the likelihood of receptor saturation or immune exhaustion is minimal. Thus, this formulation provides a biologically plausible and computationally efficient means of representing time-resolved immune input in response to multiple NP administrations.

##### Model Calibration

Experimental data for calibration comprised mean tumor burden over time for four groups: PBS, L-MSN, L-MSN-MPL, L-MSN-MPL-CpG (Figure 7). The PBS group (no immune activation)

was fit independently to estimate γ and V̶ using the logistic growth term of Equation 1 of the

model. These values were held fixed for the remaining three groups. For each treatment group, parameters **k_M_**, *k_act_*, and *d_M_* were estimated by fitting the full ODE model to group-specific mean tumor burden trajectories using nonlinear least squares. Fitting was performed on normalized data and rescaled to physical units (photons) for reporting.

##### Bootstrapping for Confidence Intervals

To quantify uncertainty, a nonparametric bootstrapping strategy was used. Tumor burden data for each group were resampled across mice to generate 200 synthetic datasets. The model was re- fit to each resampled dataset, and 95% confidence intervals (CIs) for the predicted tumor trajectories were computed from empirical quantiles.

##### Sensitivity Analysis

To evaluate the influence of model parameters (γ, V̂, **k_M_**, *k_act_*, *d_M_*, and λ) on treatment response, we performed a local sensitivity analysis using a perturbation-based approach. The analysis was conducted using the best-fit parameters obtained for the L-MSN-MPL-CpG group, which demonstrated the strongest anti-tumor response. We perturbed each model parameter individually over a symmetric range of ±50% around its best-fit value, using 101 equally spaced points. For each perturbed parameter value, we simulated the model to predict tumor burden at day 18, denoted vC18J. The initial condition for tumor burden was set to the normalized value observed in the L-MSN-MPL-CpG group at day 4, consistent with the model calibration procedure. For each parameter *P*, the sensitivity was quantified as the normalized change in tumor burden at day 18 relative to the baseline simulation:

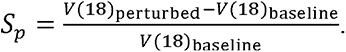

This dimensionless metric represents the relative change in model output resulting from small changes in each parameter and allows for direct comparison of parameter influence. To visualize parameter effects, we plotted the percent change in vC18J against the percent change in each parameter.

##### Predictive Simulations

To explore the potential benefits of enhanced immune activation or altered dosing regimens, we performed predictive simulations by systematically varying *k_act_* (e.g., 1.5x, 2x, 3x the fitted value) and/or including a third NP pulse. These simulations highlight how tumor burden dynamics are expected to shift under optimized nanoparticle design or scheduling.

##### Numerical Implementation

All modeling analysis and simulations were performed in MATLAB R2022b. The model was implemented using ‘ode45’ for numerical integration and ‘lsqcurvefit’ for parameter estimation.

#### 3.3.13. Statistical Analysis

Measurements in this study were obtained from distinct samples. Graphpad Prism v10.2.2 was used to perform statistical analysis. Column statistics were analyzed using an ordinary one-way ANOVA and Tukey’s multiple comparison test. For tumor burden comparisons, multiple t-tests assuming all rows were sampled from populations with the same scatter and correction for multiple comparisons using the Holm–Sidak method were used. Kaplan–Meier survival curves were analyzed using log-rank Mantel–Cox.

---

**Figure S1:**
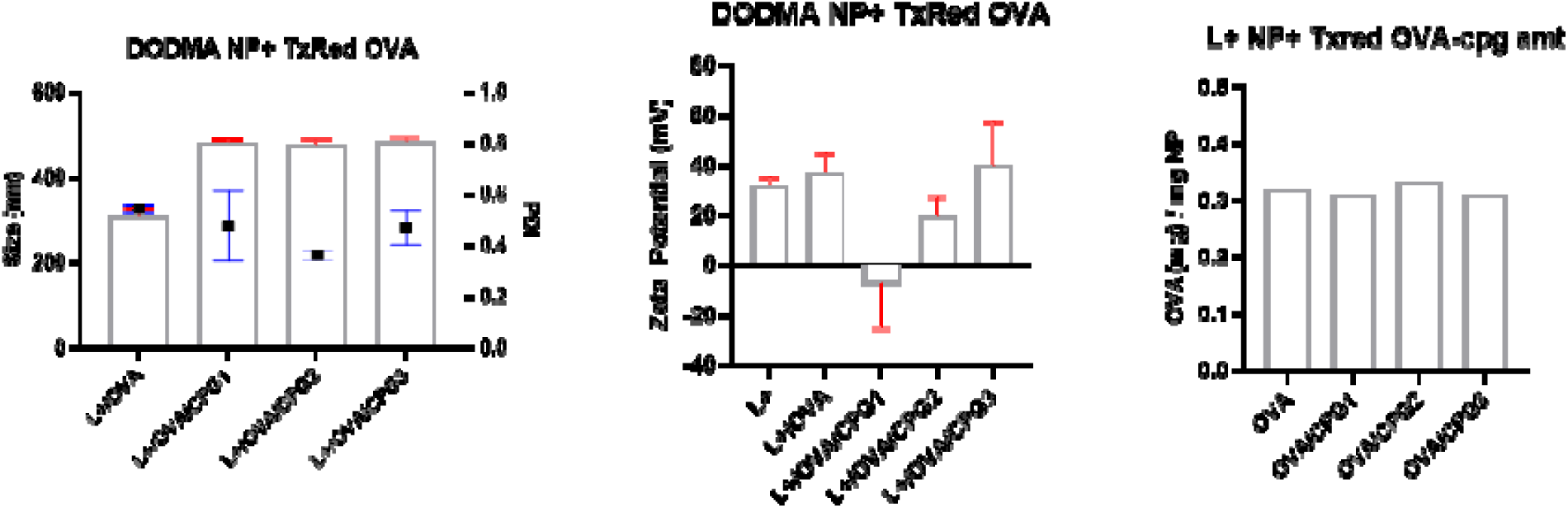
IL+ over MSN+ fail to yield homogeneous nanoparticle due to strong electrostatic repulsion between silica surface and lipids.

**Figure S2.**
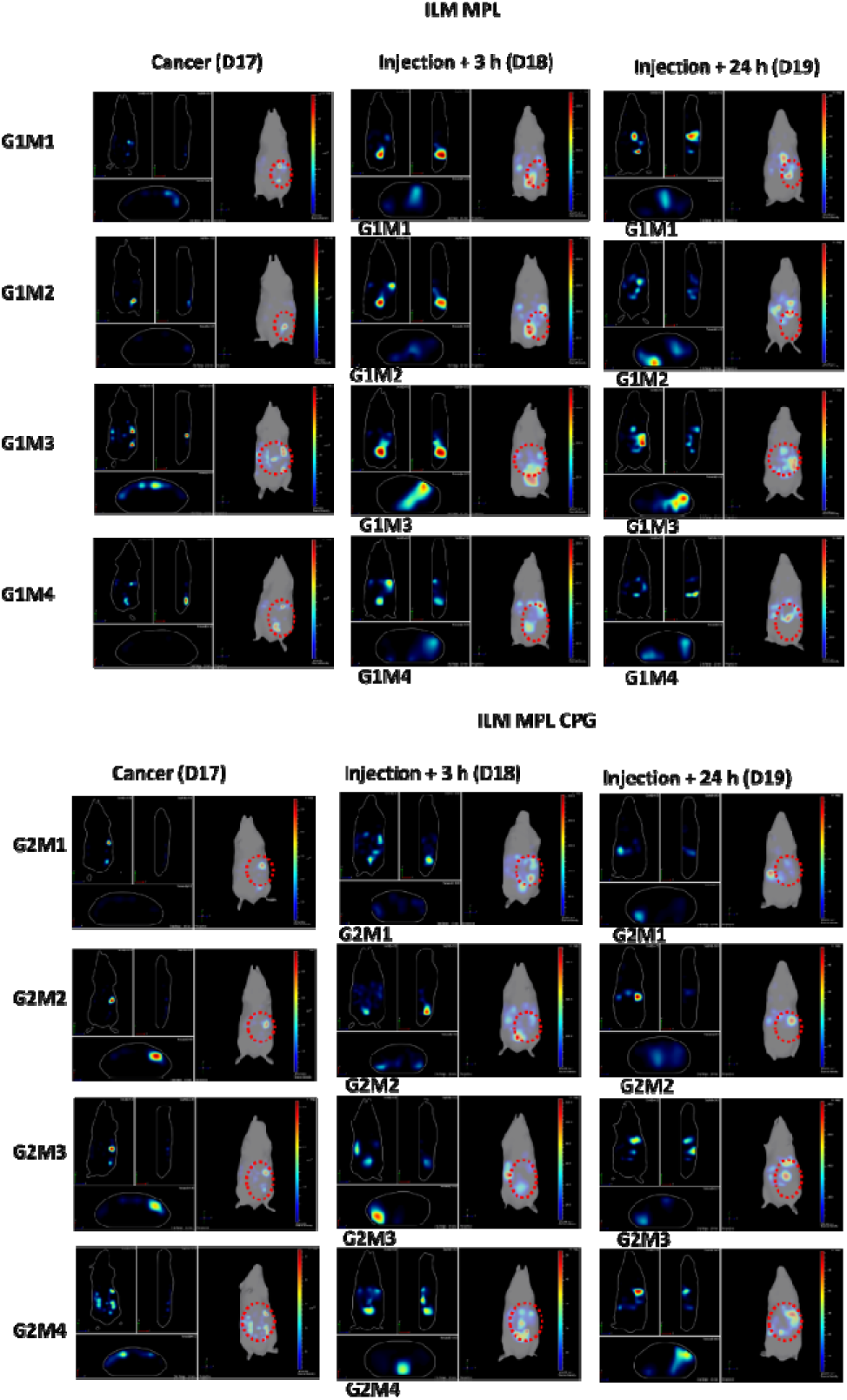
3D IVIS on living intact animal showing bioluminescent BR5 cancer cells (Day17), and Dy633-fluorescent ILM MPL and ILM MPL/CPG (Days 18 and 19)

**Figure S3.**
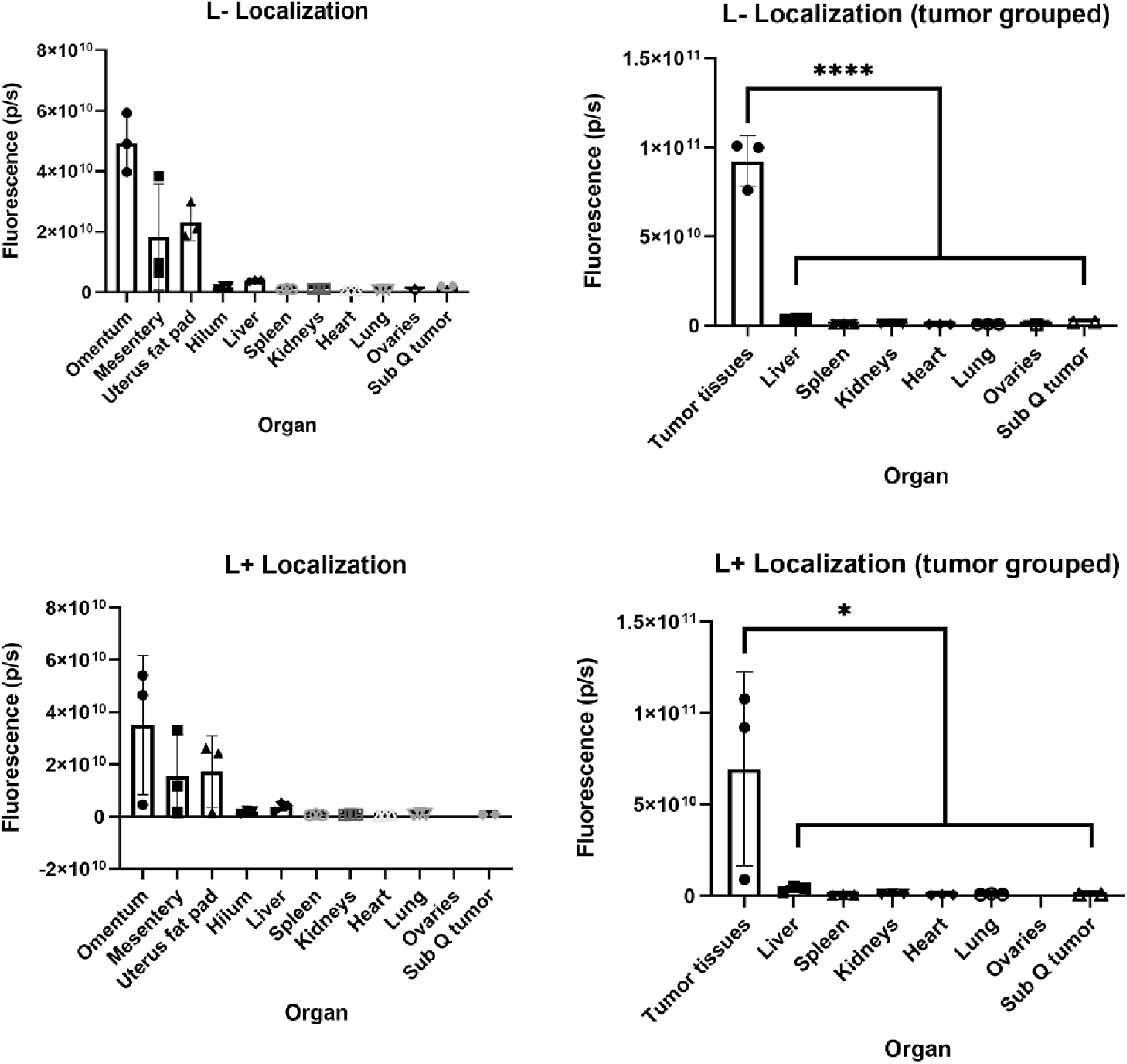
Organ accumulation of ILM- and ILM+ shows that accumulation in tumor burdened areas is charge-independent.

**Figure S4.**
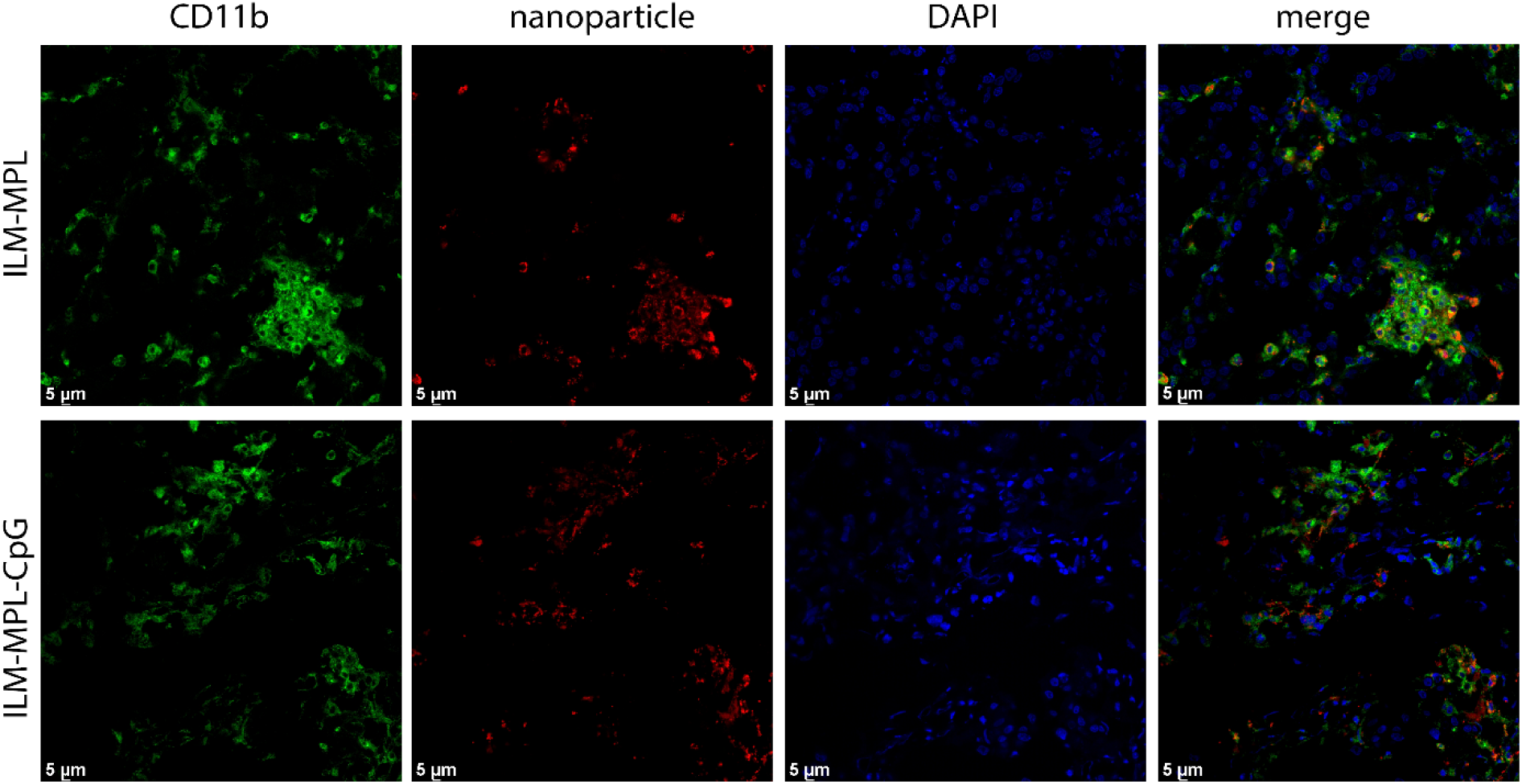
Mono or dual adjuvanted ILM (red) colocalize with CD11b (green) macrophages. Nuclei (DAPI)

**Figure S5.**
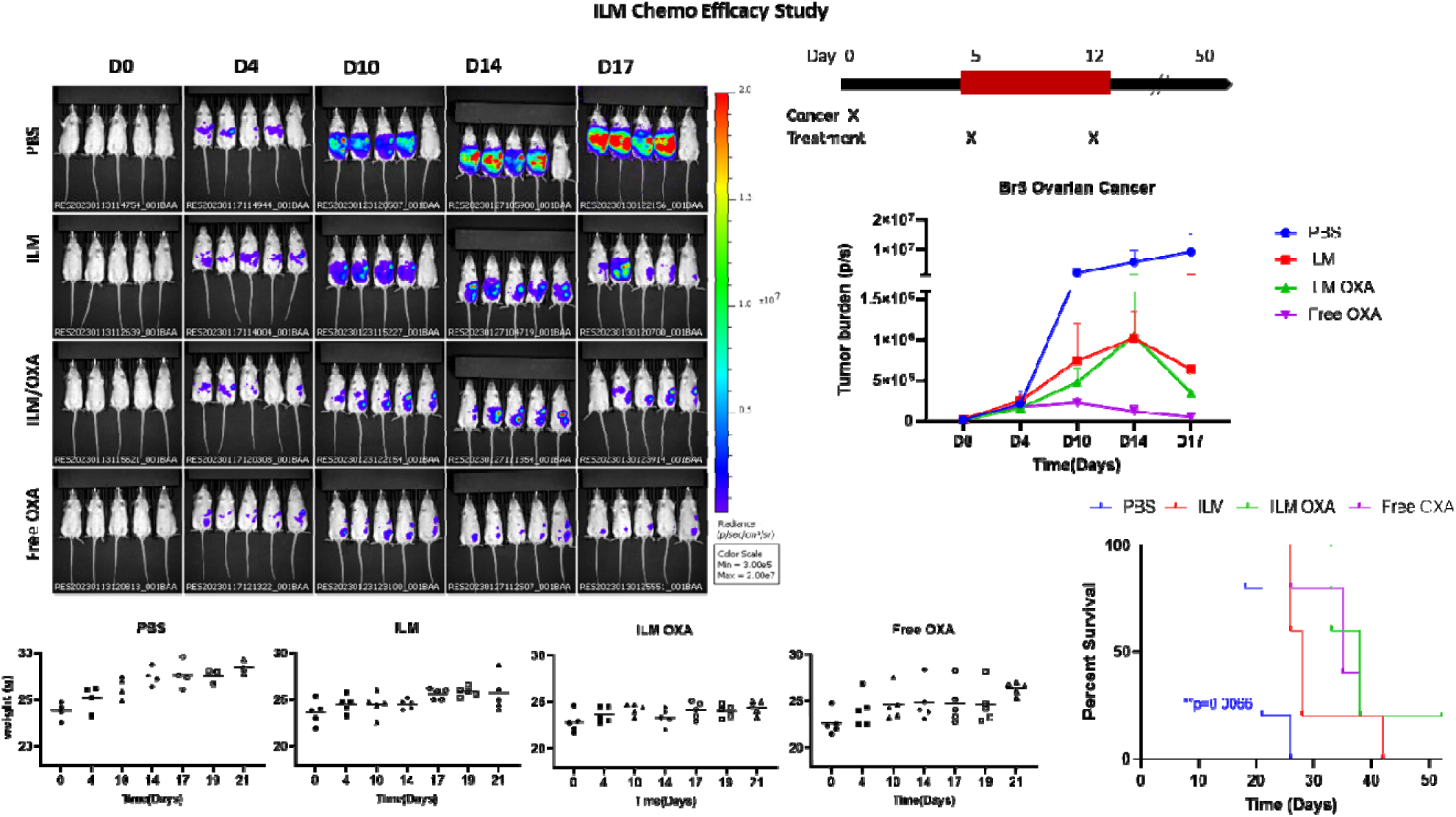
*in vivo* efficacy of oxaliplatin ILM efficacy in FVB mice

**Table S1.** The 18 liposomal formulations we have assessed in the current study. Numbers are mol% of each lipid within the IL formulation. For consistency, we indexed all lipids as IL. We underlined formulations are the selected for the negative and positive ILM.

| IL # | DPPC | DOPC | DOPE | EGGSM | Cholesterol | DMPG | DOTAP | DSTAP | DODMA | MPL |
| --- | --- | --- | --- | --- | --- | --- | --- | --- | --- | --- |
| **Negative ILs (no use of positive lipids)** |  |  |  |  |  |  |  |  |  |  |
| **1** | 99.5 | - | - | - | - | - | - | - | - | 0.5 |
| **2** | 96.5 | - | - | - | 3 | - | - | - | - | 0.5 |
| **3** | 74.5 | 5 | - | - | 20 | - | - | - | - | 0.5 |
| **4** | 74.5 | - | 5 | - | 20 | - | - | - | - | 0.5 |
| **5** | 29.5 | - | - | - | 10 | 50 | - | - | - | 0.5 |
| **6** | 50 | - | - | - | 10 | 40 | - | - | - | - |
| **7** | 49.5 | - | - | - | 10 | 40 | - | - | - | 0.5 |
| **8** | 48 | - | - | - | 10 | 40 | - | - | - | 2.0 |
| **Positive ILs (use of positive lipids)** |  |  |  |  |  |  |  |  |  |  |
| **9** | 65 | - | - | - | 25 | - | 10 | - | - | - |
| **10** | 64.5 | - | - | - | 25 | - | 10 | - | - | 0.5 |
| **11** | 60 | - | 20 | - | 10 | - | 10 | - | - | - |
| **12** | 70 | - | 10 | - | 10 | - | 10 | - | - | - |
| **13** | - | - | - | 80 | 10 | - | 10 | - | - | - |
| **14** | 60 | - | - | 20 | 10 | - | 10 | - | - | - |
| **15** | 70 | - | 10 | - | 10 | - | - | 10 | - | - |
| **16** | 68 | - | 20 | - | 10 | - | - | - | - | 2 |
| **17** | 76 | - | - | - | 13 | - | - | - | 11 | - |
| **18** | 74 | - | - | - | 13 | - | - | - | 11 | 2 |

**Table S2.** quality control assessment of the ILs and corresponding ILMs made. The formulations were first screened as a function of hydrodynamic size and zeta potential as after fusion onto MSN (ILM). The cut-off hydrodynamic size was ∼ 250 nm whilst PDI’s was ∼ 0.25 for ILM and 0.3 for ILs. Light colors indicate promising formulations, dark red indicates dropped formulations and dark green indicate the lipid formulations we have used in this study.

| Immunogenic Liposomes (IL) |  |  |  | Immunogenic Lipid-Coated MSNs (ILM) |  |  |
| --- | --- | --- | --- | --- | --- | --- |
| | Size (std dev) (nm) | PDI (± std dev) | Zeta Potential (± std dev) (mV) | Size (± std dev) (nm) | PDI (± std dev) | Zeta Potential (± std dev) (mV) |
| **Theoretically Negative ILMs (negative ILs fused on positive MSN)** |  |  |  |  |  |  |
| **1** | 194.3 (17.3) | 0.59 (0.05) | -4.8 (0.73) | 213.5 (7.9) | 0.31 (0.04) | -19.9 (3.16) |
| **2** | 43.8 (0.4) | 0.26 (0.01) | -4.5 (0.85) | 259.4 (4.8) | 0.35 (0.02) | -17.2 (2.40) |
| **3** | 62.7 (2.2) | 0.32 (0.04) | -8.9 (0.25) | 820.9 (125.5) | 0.25 (0.07) | -17.8 (0.95) |
| **4** | 55.1 (0.5) | 0.33 (0.01) | -7.3 (0.71) | 233.2 (15.9) | 0.81 (0.13) | -9.1 (5.10) |
| **5** | 31.0 (2.2) | 0.09 (0.01) | -55.1 (2.93) | 342.4 (3.4) | 0.45 (0.03) | -33.3 (2.41) |
| **6** | 24.9 (1.6) | 0.15 (0.01) | -54.8 (0.7) | 208.8 (1.5) | 0.21 (0.01) | -27.8 (1.14) |
| **7** | 23.8 (1.8) | 0.18 (0.02) | -55.1 (1.67) | 229.8 (6.4) | 0.17 (0.03) | -27.8 (1.56) |
| **8** | 27.0 (1.4) | 0.31 (0.01) | -56.1 (5.28) | 230.6 (3.8) | 0.14 (0.02) | -27.9 (0.49) |
| **Theoretically Positive ILMs (Positive ILs fused on negative MSN)** |  |  |  |  |  |  |
| **9** | 38.7 (3.3) | 0.20 (0.01) | 42.6 (2.52) | 297.8 (7.8) | 0.21 (0.05) | 8.8 (0.65) |
| **10** | 55.1 (1.3) | 0.22 (0.01) | 47.6 (3.24) | 569.3 (76.1) | 0.20 (0.17) | 3.0 (1.27) |
| **11** | 41.1 (0.2) | 0.19 (0.006) | 9.5 (3.3) | 300.5 (6.23) | 0.17 (0.004) | 12.5 (0.36) |
| **12** | 50.6 (0.09) | 0.25 (0.007) | 8.1 (3.2) | 580 (8.3) | 0.23 (0.02) | 23.7 (1.76) |
| **13** | 42.5 (0.35) | 0.25 (0.01) | 15.5 (1.3) | 587.7 (18.6) | 0.27 (0.02) | 14.4 (1.11) |
| **14** | 55.5 (0.7) | 0.25 (0.008) | 18.6 (4.8) | 568.6 (6.8) | 0.40 (0.01) | 25.2 (0.25) |
| **15** | 42.1 (0.7) | 0.22 (0.01) | 26.4 (1.1) | 402.4 (2.4) | 0.26 (0.03) | 22.8 (0.49) |
| **16** | 58.7 (0.8) | 0.26 (0.01) | -7.8 (0.61) | 896.4 (21.5) | 0.25 (0.02) | -12.3 (1.6) |
| **17** | 90.8 (2.2) | 0.28 (0.03) | 31.9 (3.2) | 242.7 (3.25) | 0.24 (0.004) | 6.93 (8.8) |
| **18** | 91.2 (0.9) | 0.27 (0.03) | 10.2 | 215.6 (9.2) | 0.20 (0.005) | 8.92 (0.46) |

**Table S3.**
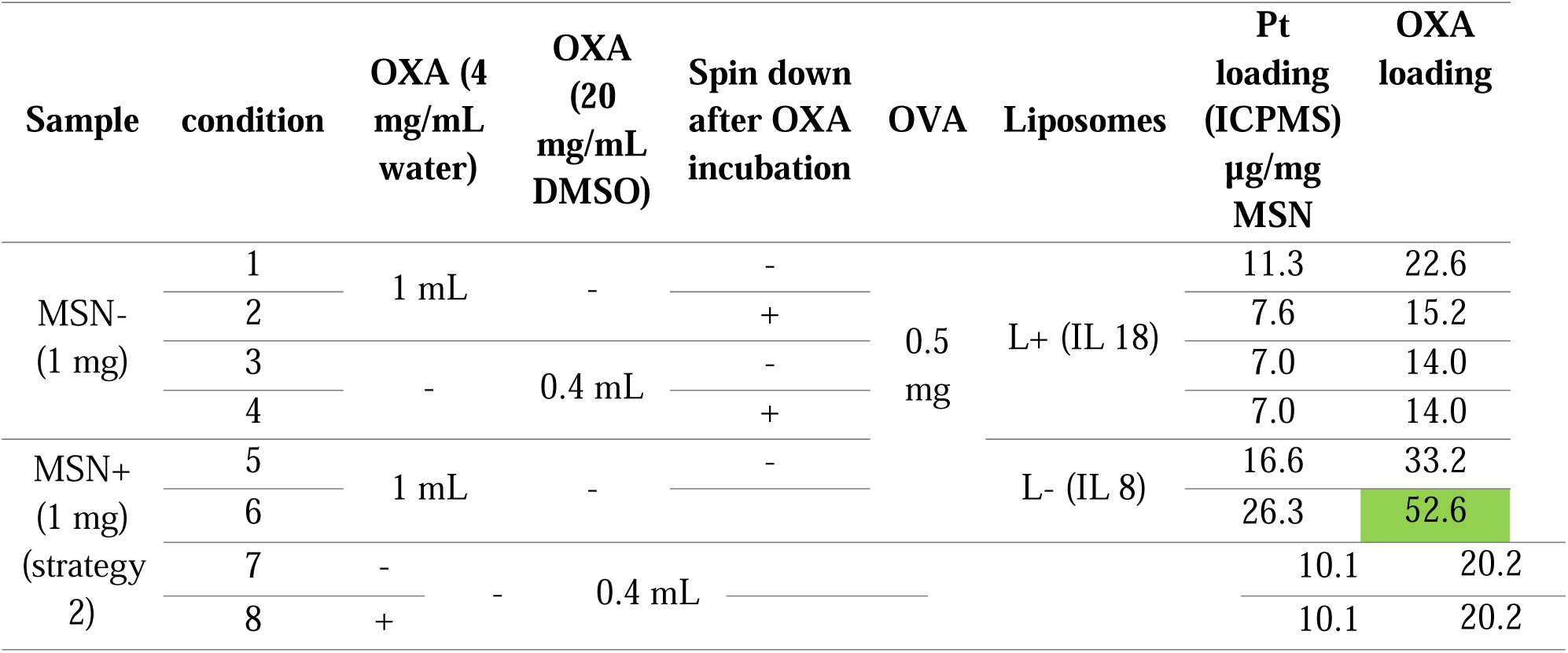
Oxaliplatin loading and quantification in ILM- (MSN+) and ILM+ (MSN-). Strategies used for OXA loading on ILM8 and ILM18 and OXA amount per mg MSN measured by ICPMS.

**Table S4.** List of mathematical model parameters.

| Parameter | Definition | Units | Value | 95% CI |
| --- | --- | --- | --- | --- |
| $\gamma$ | Tumor growth rate constant | 1/day | 0.468 | [0.399 - 0.550] |
| $\hat{V}$ | Tumor carrying capacity | Photons | 1.79e+9 | [1.08e+9 – 2.78e+9] |
| $k_M$ | Immune-mediated tumor killing rate | 1/day·photons | Varies by group | See below |
| $k_{\text{act}}$ | NP-induced immune activation rate | 1/day | Varies by group | See below |
| $d_M$ | Decay rate of immune effector cells | 1/day | Varies by group | See below |
| $\lambda$ | Immune stimulation decay rate (NP signal) | 1/day | $\log(2)/3$ | - |

**Fitted Parameters per Treatment Group**
| Group | $k_M$ [1/day·photons] | $k_{\text{act}}$ [1/day] | $d_M$ [1/day] |
| --- | --- | --- | --- |
| L-MSN | 4.19e-9 [4.00e-18–7.85e-8] | 0.81 [0.15–13.75] | 0.74 [0.18–14.82] |
| L-MSN-MPL | 5.86e-9 [4.28e-10–1.28e-8] | 0.56 [0.49–7.78] | 0.0025 [1e-4–0.285] |
| L-MSN-MPL-CpG | 5.47e-9 [3.9e-10–1.3e-8] | 0.77 [0.50–12.06] | 0.0001 [1e-5–0.19] |

## Notes

### Competing Interest Statement

The authors have declared no competing interest.

